# Minimal biophysical model of combined antibiotic action

**DOI:** 10.1101/2020.04.18.047886

**Authors:** Bor Kavčič, Gašper Tkačik, Tobias Bollenbach

**Affiliations:** Institute of Science and Technology Austria, Am Campus 1, A-3400 Klosterneuburg, Austria; Institute for Biological Physics, University of Cologne, Zülpicher Str. 77, D-50937 Cologne, Germany

## Abstract

Phenomenological relations such as Ohm’s or Fourier’s law have a venerable history in physics, but are still scarce in biology. This situation restrains predictive theory. Here, we build on bacterial “growth laws,” which capture physiological feedback between translation and cell growth, to construct a minimal biophysical model for the combined action of ribosome-targeting antibiotics. Our model predicts drug interactions like antagonism or synergy solely from responses to individual drugs. We systematically refine the model by including direct physical interactions of different drugs on the ribosome. In a limiting case, our model provides a mechanistic underpinning for recent predictions of higher-order interactions derived using entropy maximization. It further makes parameter-free predictions for combined drug effects on cells carrying resistance genes and for drugs that mimic poor nutrient environments. We show experimentally that resistance genes can drastically alter drug interactions in notable agreement with our theoretical predictions. While minimal, the model is readily adaptable and opens the door to predicting interactions of second and higher-order in a broad range of biological systems.

## I. INTRODUCTION

Antibiotics are small molecules that interfere with essential processes in bacterial cells, thereby inhibiting growth or even killing bacteria [1]. Even though antibiotics have been used in the clinic for nearly a century and have molecular targets that are often known, bacterial responses to antibiotics are complex and largely unpredictable. Due to the looming antibiotic-resistance crisis [2], understanding and predicting bacterial responses to antibiotics is becoming increasingly important. A promising way to fight the emergence and spread of resistant pathogens is to use more than one drug simultaneously [3]; however, predicting the effects of such drug combinations is a great challenge.

The combined effect of antibiotics emerges from the complex interplay of individual drug effects and the physiological response of the cell to the drug combination. Drug interactions are determined by the combined effect of multiple drugs on cell growth and survival. These interactions are defined with respect to an additive reference. By definition, additive drugs act as substitutes for each other; synergy occurs if the combined effect of the drugs is stronger than in the additive reference case, and antagonism occurs if the combined effect is weaker. An extreme case of antagonism– termed suppression–occurs when one of the drugs loses its potency in the presence of the other drug, *i.e*., bacterial growth is accelerated by adding one of the drugs. In practice, such drug interactions are determined by measuring bacterial growth in a two-dimensional assay in which each drug is dosed in a gradient along each axis.

By measuring the growth rate λ over a two-dimensional matrix of drug concentrations (*c_A_, c_B_*), the dose-response surface *y*(*c_A_, c_B_*) = λ(*c_A_, c_B_*)/λ_0_ is obtained; here, λ_0_ is the growth rate in the absence of antibiotics. The doseresponse surface can be characterized by the shape of its contours, *i.e*., the lines of equal growth rate (isoboles). By definition, in the additive case these contours are linear (but not necessarily parallel) (Fig. 1). Linear isoboles imply that increasing the concentration of one drug is equivalent to increasing the concentration of the other. Drug interactions according to the Loewe definition [4] occur when the dose-response surface deviates from this additive expectation. The interaction is synergistic or antagonistic if the drug combination is more or less potent than the additive expectation, respectively (Fig. 1). There are other definitions of drug interactions: In the Bliss definition, the responses to individual drugs are multiplied to yield the reference response to the combination [5, 6]. In this work, we use the Loewe definition, but we are principally interested in the shape of entire dose-response surfaces, which is independent of the exact definition of drug interactions.

**FIG. 1.**
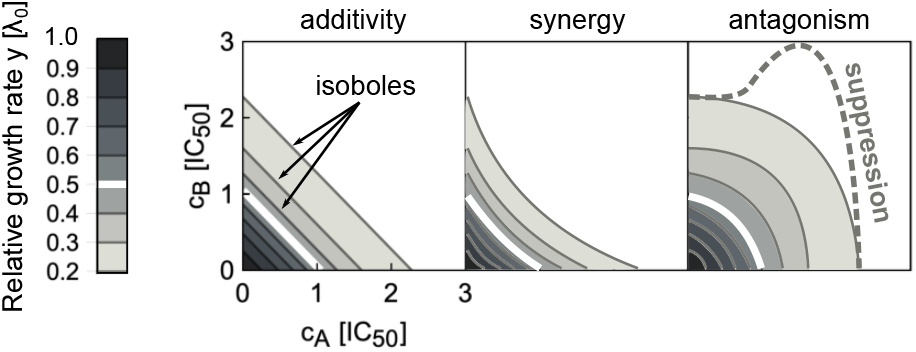
Drug interaction types are defined by the shape of the dose-response surface. The dose-response surface is given by the relative growth rate *y* as a function of the concentrations of two drugs (*c_A_, c_B_*). Here, the concentrations are normalized such that *c_i_* = 1 corresponds to 50% growth inhibition in the presence of drug *i* alone. The dose-response surface is defined by lines of constant growth rate (isoboles); the white isobole corresponds to 50% growth inhibition; different shades of gray correspond to different relative growth rates. Additivity (left) represents the case in which antibiotics act as substitutes for each other. Synergy (middle) and antagonism (right) are characterized by convex or concave isoboles (curving towards or away from the origin), respectively. Isoboles corresponding to suppression (dashed gray) are non-monotonic; in the example shown, this implies that adding drug A on top of drug B increases the growth rate.

Higher-order interactions, which occur when more than two drugs are combined, can be predicted to some extent using mechanism-independent models [7, 8]. However, such predictions generally require prior knowledge of the pairwise interactions between all drugs involved, which can usually not be predicted. One of the main reasons for this situation is that the underlying mechanisms of drug interactions are largely unknown. Systematic measurements of all pairwise drug interactions are hampered by the combinatorial explosion of possible drug-drug and concentration-concentration combinations, which prohibit such brute-force approaches. Thus, to guide the design and analysis of drug combinations, it is crucial to develop predictive theoretical models.

Apart from their clinical importance, antibiotics targeting the bacterial ribosome (translation inhibitors) are particularly well-suited for biophysical modeling since the physiological response to perturbations of translation can be described quantitatively using bacterial growth laws [9, 10]. These empirical relations offer a phenomenological description of the growth-dependent state of the bacterial cell and provide a solid foundation for quantitative studies of bacterial physiology. Similar to laws in physics, such as Fourier’s law of heat conduction or Ohm’s law, these phenomenological relations enable the construction of predictive mathematical models without free parameters even if their microscopic origins are not yet understood [11]. Translation inhibitors have the additional advantage that many of the drug interactions occurring between them have been recently measured [12].

Here, we present a biophysical model that predicts bacterial growth responses to combinations of translation inhibitors. Starting from responses to single antibiotics, we derive approximate analytical solutions of this model and investigate the effects of direct physical or allosteric interactions between antibiotics on the ribosome. We discuss several relevant extensions of the model, in particular (1) interactions with antibiotics that induce starvation, (2) the effects of resistance genes, (3) the correspondence to non-mechanistic models of interactions between more than two drugs, and (4) predictions for interactions of translation inhibitors with antibiotics that alter growth law parameters. We validate several non-trivial predictions made by the biophysical model in experiments.

## II. MODEL FOR A SINGLE TRANSLATION INHIBITOR

First, we recapitulate the biophysical model for a single translation inhibitor [10]. The model captures the kinetics of antibiotic transport into the cell and binding to the ribosome (Fig. 2a,b), as well as the physiological response of the cell to translation perturbation. This physiological response is described by bacterial growth laws, which summarize the interdependence of the intracellular ribosome concentration *r* and the growth rate λ (Fig. 2b). Bacterial physiology and the response to antibiotic treatment strongly depend on the nutrient environment. In particular, the number of ribosomes per cell varies over approximately 5 – 75 × 10^3^ [13]. The ribosome concentration increases linearly with the growth rate when the latter is varied by changing the quality of the growth medium:

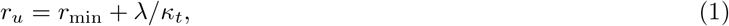

where *κ_t_* = 0.06 *μ*M^−1^h^−1^, *r_u_* and *r*_min_ = 19.3 *μ*M are the translational capacity, the concentration of unperturbed ribosomes, and a minimal ribosome concentration, respectively [9, 10]. This first growth law states that unperturbed ribosomes synthesize new proteins, whose overall synthesis rate is proportional to the growth rate. This relation holds across diverse growth media and different *Escherichia coli* strains [9]. Typical values for doubling times range from hours to approximately twenty minutes, corresponding to growth rates up to around 2.5 h^−1^. However, when the growth rate is lowered by addition of a translation inhibitor in a constant nutrient environment, the total ribosome concentration *r*_tot_ and growth rate become negatively correlated and obey:

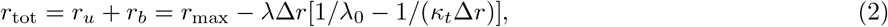

where *r*_max_ = 65.8 *μ*M, Δ*r* = *r*_max_ – *r*_min_ = 46.5 *μ*M and λ_0_ are the maximal ribosome concentration, the dynamic range of ribosome concentration, and the growth rate in the absence of antibiotics, respectively [9, 10]. Eq. (2) quantitatively describes the upregulation of ribosome production that occurs in response to translation inhibition: Bacteria produce more ribosomes to compensate for the ribosomes blocked by antibiotics.

**FIG. 2.**
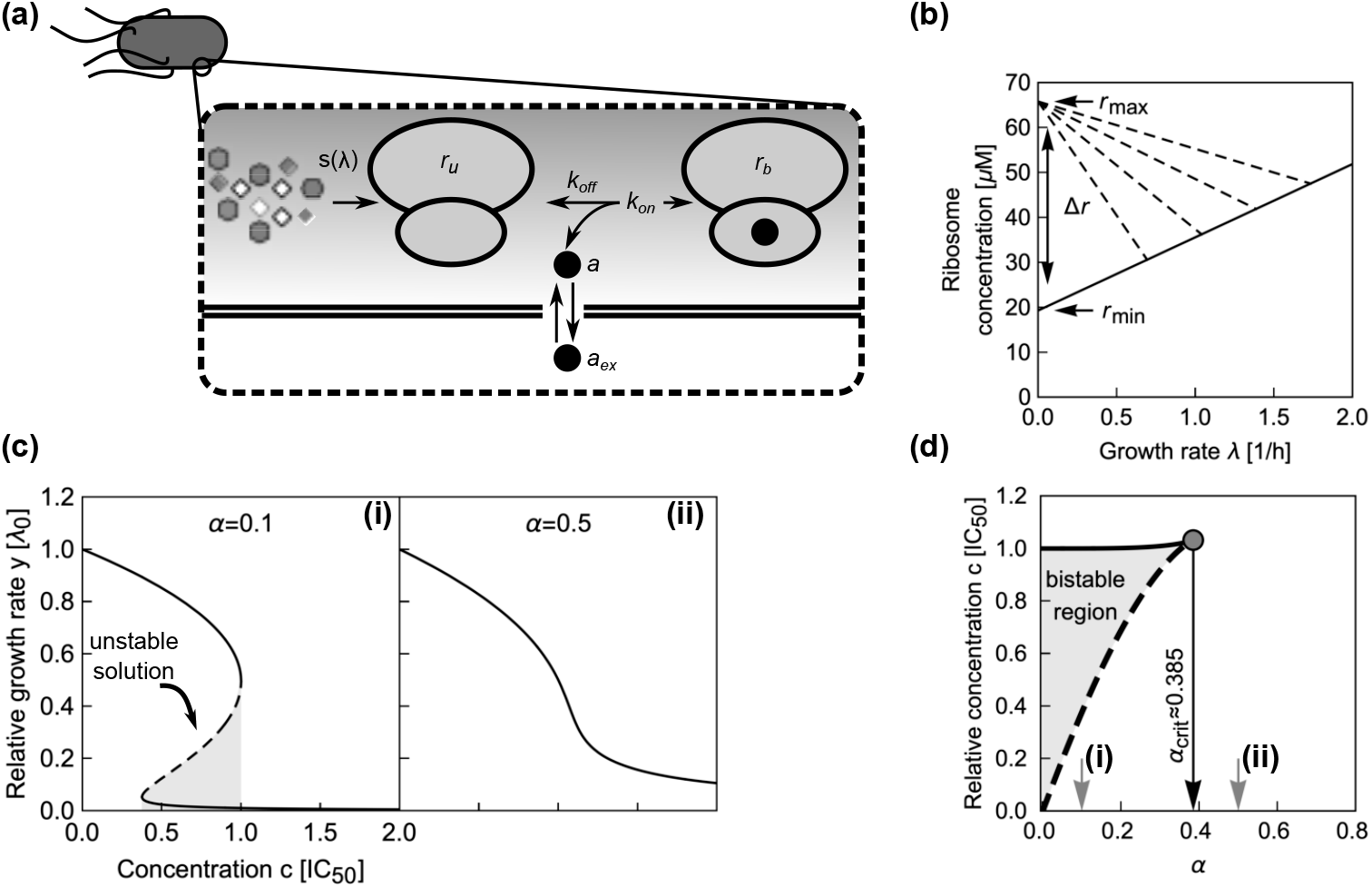
Main components of the model for a single translation inhibitor and exemplary dose-response curves. (a) Schematic of processes captured by the model. Ribosomes (double ovals) are synthesized with the rate *s*(λ) and are initially unbound by antibiotic (*r_u_*). Unbound ribosomes contribute to growth. Antibiotics enter the cell (*a*_ex_ → *a*) and bind to and detach from ribosomes with second-order and first-order rate constants *k*_on_ and *k*_off_, respectively. Bound ribosomes (*r_b_*) do not contribute to growth. (b) Bacterial growth laws. When the growth rate is varied by changing the quality of the nutrient environment, the ribosome concentration increases linearly with growth rate (solid line). If growth is inhibited by a translation inhibitor, the ribosome concentration increases with decreasing growth rate (dashed lines). The intercepts of the solid and dashed lines determine the minimal (*r*_min_) and maximal ribosome concentration (*r*_max_), respectively, which are Δ*r* apart. (c) Examples of dose-response curves. The model can produce both (i) steep and (ii) shallow dose-response curves, depending on the parameter *α* (see text). The steep dose-response curve has a region of concentrations (gray shaded area) where one unstable (dashed line) and two stable (solid lines) solutions exist. (d) Exact phase diagram for dose-response curves. The shaded area shows the region of drug concentrations where two stable solutions exist. Gray arrows show *α* for the examples from (c); black arrow shows the critical value 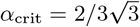 above which no bistability can occur.

When antibiotics enter the cell, they can bind to ribosomes. The net rate of forward and reverse binding of antibiotics to the ribosome is given by *f*(*r_u_, r_b_, a*) = −*k*_on_*a*(*r_u_* – *r*_min_) + *k*_off_*r_b_*, where *k*_off_ and *k*_on_ are first and second order rate constants, and *a* is the intracellular antibiotic concentration (Fig. 2a). Here, we assumed that only ribosomes capable of translation (*r*_u_ – *r*_min_) can be bound by the antibiotics [10, 14].

The intracellular antibiotic concentration is affected by the kinetics of antibiotic entry into the cell, which is given by *J*(*a*_ex_, *a*) = *p*_in_*a*_ex_ – *p*_out_*a*, where *a*_ex_ is the extracellular antibiotic concentration. Typical influx and efflux rates, *p*_in_ and *p*_out_, for different translation inhibitors range from 1 – 1000 h^−1^ and from 0.01 – 100 h^−1^, respectively. Typical rates of forward and reverse binding, *k*_on_ and *k*_off_, are around 1000 *μ*M^−1^h^−1^ and between 0 – 10^5^ h^−1^, respectively [10, 14]. Here, *k*_off_ = 0 corresponds to antibiotics with effectively irreversible binding such as streptomycin [10, 15]. All molecular species in the cell are effectively diluted at rate λ as cells grow and divide. Since the ribosome concentration is determined by Eq. (2), the ribosome synthesis rate *s* depends on the growth rate, *i.e., s* = *s*(λ). Together, these terms constitute a closed system of ordinary differential equations:

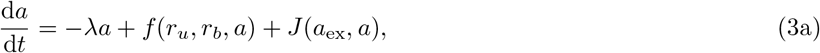

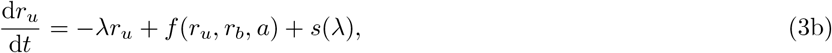

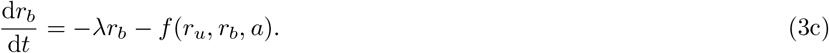

Here, the ribosome synthesis rate reads *s*(λ) = λ*r*_tot_ = λ {*r*_max_ – λΔ*r*[1/λ_0_ – 1/(*κ_t_*Δ*r*)]}. The steady-state solution of Eqs. (3) represents a balanced-growth state of the system – the situation that is commonly investigated in experiments.

The steady-state solution reads [10]

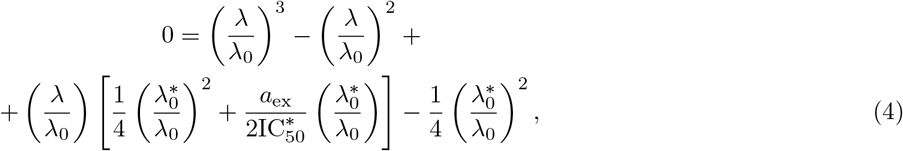

where 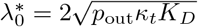 with *K_D_* = *k*_off_/*k*_on_, and 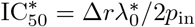. We can recast Eq. (4) into (Appendix A):

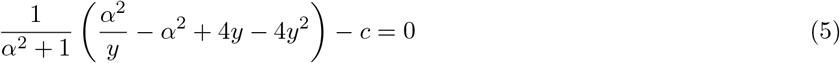

by defining *a*_ex_ = *c* × IC_50_, λ = *y* × λ_0_ and 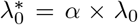, where IC_50_ is the extracellular antibiotic concentration that leads to 50% growth inhibition, a common measure of drug sensitivity. Here, we call *α* the response parameter, as it describes the dose-response curve shape: The higher the value taken by *α*, the shallower the dose-response curve (Fig. 2c).

Since Eq. (5) is cubic in the relative growth rate *y*, there are generally either one or three real solutions for *y* (Fig. 2c). This indicates that there is a parameter regime in which the dynamical system can exhibit bistability [10, 16]. Previous studies identified the bistable parameter regions numerically or in closed expression with many parameters [10, 14]. Notably, the rescaling shown above enables the exact calculation of the bifurcation point (see Appendix A): When 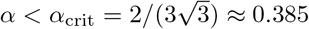 the system can be bistable (Fig. 2d), *i.e*., there is a region of concentrations with stable solutions at two different growth rates. In this region, the growth rate sharply declines when a critical concentration is exceeded. It is difficult to measure such steep dose-response curves experimentally since very low growth rates are challenging to detect and quantify. Additionally, bistability cannot be observed in population-level experiments since the high-growth-rate branch will quickly dominate the population; single-cell experiments are needed to observe growth bistability [17]. On the other hand, if the antibiotic concentration can be varied during the experiment, bistability can be tested by determining the hysteresis of the response, as observed for synthetic gene networks [18].

Steep dose-response curves (*α* < *α*_crit_) occur for antibiotics with tight binding to the ribosome (*K_D_* → 0) or inefficient efflux (*p*_out_ → 0). Alternatively, if these two quantities are growth-rate invariant, dose-response curves become steeper with increasing growth rate in the absence of drug, as *α* ∝ 1/λ_0_. For typical values of the relevant parameters (discussed above), *α* ranges from 0 to ~ 10. We have experimentally observed values of *α* for different translation inhibitors in the range 0 – 2 [12]. In the limit *α* ≫ 1, Eq. (5) simplifies into *y* = 1/ (1 + *c*); if *α* → 0 then this expression becomes 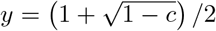 for *c* ≥ 1, [10]. This biophysical model for a single translation inhibitor provides the foundation for a predictive theory of multiple drug interactions between different translation inhibitors.

## III. MODEL FOR INTERACTION BETWEEN TWO TRANSLATION INHIBITORS

### A. Dynamical system describing the binding of antibiotics and the physiological state of the cell

When combinations of two different translation inhibitors are present, each ribosome can be bound by either of them alone or by both simultaneously. To generalize the model described in Sec. II to this situation, we need to introduce additional populations of ribosomes. Extending the mathematical model [Eqs. (3)] to two translation inhibitors yields:

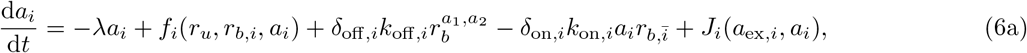

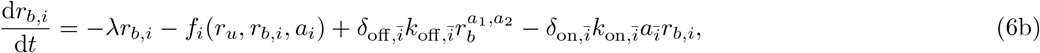

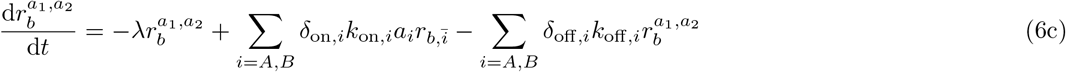

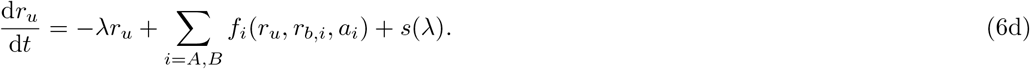

The terms *f*_i_(*r*_u_, *r_b,i_*, *a_i_*) and *J_i_*(*a*_ex,*i*_, *a_i_*) describe the first binding step and membrane transport of antibiotic *i*, respectively. The additional terms 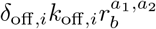 and 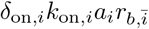 describe the unbinding of antibiotic *i* from double-bound ribosomes 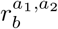 and the binding of antibiotic 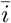 to ribosomes already bound by the other antibiotic (*e.g*., for antibiotics *A* and *B*, *Ā* = *B*), respectively. The dimensionless parameters *δ_σ,i_* with *σ* ∈ {on, off} denote the relative change of the rate of forward and reverse binding of antibiotic *i* to ribosomes already bound by the other antibiotic. When the binding kinetics for both antibiotics are independent, all *δ_σ,i_* = 1. When both antibiotics compete for the same binding site on the ribosome, *δ*_on,*i*_ = 0. In general, the parameters can vary continuously to capture any changes in ribosome binding of one antibiotic due to the binding of the second.

What is the main consequence of including the double-bound ribosomes? Below, we show that in the absence of double-bound ribosomes, drug interactions are generally expected to be additive. If we assume that no double-bound ribosomes can form, *e.g*., by setting *δ*_on,*i*_ = 0, Eq. (6c) becomes equal to zero and all terms associated with the second binding event disappear. To show that this situation necessarily yields an additive drug interaction, we examine the shape of the isoboles. Along an isobole, *i.e*., at fixed growth rate, *r_u_* = λ/*κ_t_* + *r*_min_ is constant. This implies that the total concentration of ribosomes bound by one antibiotic remains constant for all different pairs (*c_A_, c_B_*) along the isobole. In steady state, the concentration of ribosomes bound by antibiotic *i* is *r_b,i_* = *a_i_* × *k*_on,*i*_λ/ [(*k*_off,*i*_ + λ) *κ_t_*] = *a_i_* × *ξ_i_*. Thus, the sum of single-bound ribosome concentrations is

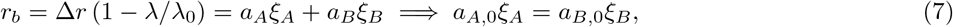

where *a*_*A*,0_ and *a*_*B*,0_ are the intracellular concentrations of antibiotics *A* and *B* in the absence of the other antibiotic, respectively. We express *a_i_* as a function of *a*_ex,*i*_ from Eq. (6a), which yields:

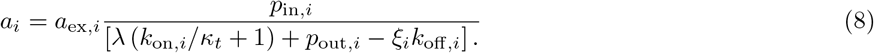

The proportionality constant in this expression depends only on λ and kinetic parameters; in particular, it is invariant of the concentration of the other antibiotic. Thus, Eq. (7) corresponds to a linear isobole. This argument shows that additivity generally occurs when double-bound ribosomes cannot form. Setting *δ*_on,*i*_ = 0 is a convenient way to obtain an additive expectation from the model, which is useful as a reference for the case in which double-bound ribosomes can form.

To study the effect of double-bound ribosomes (Fig. 3a), we systematically calculated dose-response surfaces for both competitive and independent binding (Fig. 3b; Appendix C). The Loewe interaction score (*LI*) is a convenient way to characterize the type and strength of drug interactions by a single number, with negative values corresponding to synergy and positive values to antagonism [12]. The *LI* score quantifies the interaction using the volume under the dose-response surface:

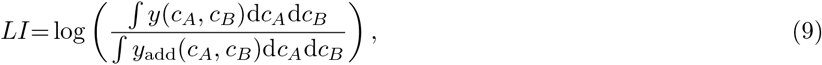

where *y*_add_ is the response surface of the additive expectation, which is calculated directly from the responses to the individual drugs (see Appendix B). By calculating the *LI* score of the dose-response surfaces for varying response parameters *α_A_, α_B_* of the two antibiotics that are combined, we determined the complete phase diagram of drug interactions (Fig. 3c). This procedure revealed that antagonism generally occurs for combinations of antibiotics with steep dose-response curves, and the interaction becomes additive and then synergistic with increasing response parameters (Fig. 3b). This transition from antagonism to synergy is smooth and partitions the phase diagram into two regions (Fig. 3c). We can directly use this phase diagram to test if experimentally observed drug interactions [12] agree with those calculated based on the measured response parameters *α* of the individual drugs. Indeed, additive and mildly antagonistic interactions produced by this theoretical description agree with experimental observations [12]; two examples are shown in Fig. 3c. However, suppressive interactions remain elusive.

**FIG. 3.**
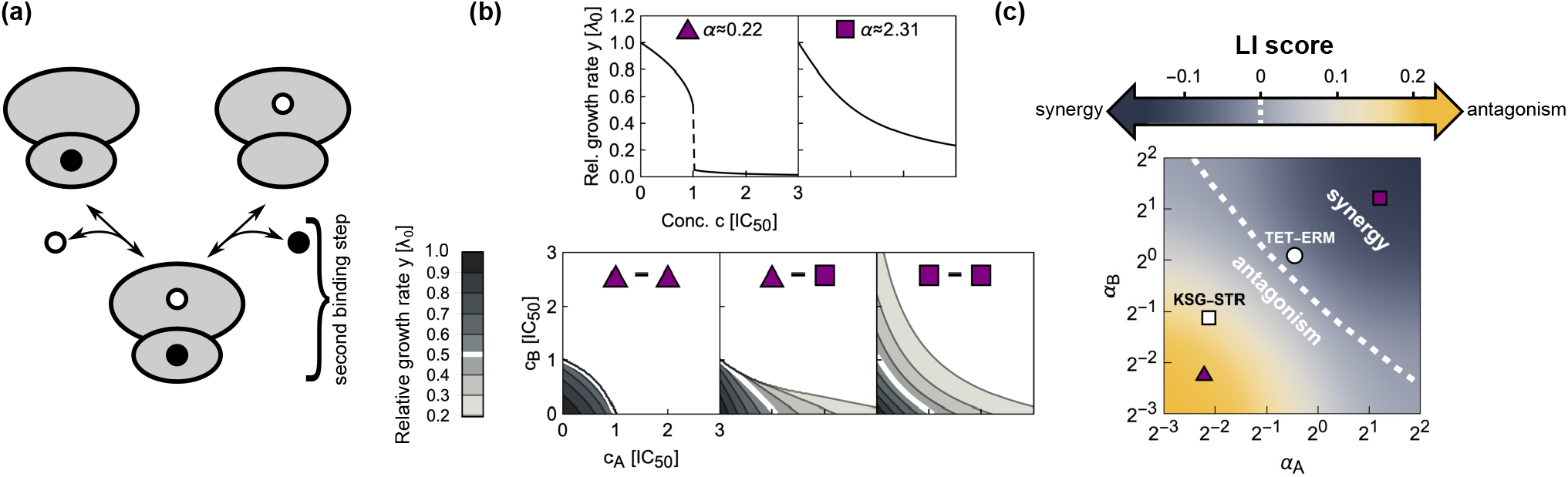
Biophysical model of two antibiotics that can bind the ribosome simultaneously produces different types of drug interaction. (a) Schematic: Ribosomes already bound by a single antibiotic (black and white circles) can be bound by another one. If the binding is independent of the presence of an already bound antibiotic, the second binding step follows the same kinetics as for a single antibiotic. (b) Examples of dose-response curves of different steepness and corresponding dose-response surfaces calculated from the model. Top: Dose-response curves with low or high *α* are steep (left) or shallow (right), respectively. Bottom: Depending on the shape of the dose-response curves of the antibiotics that are combined, the calculated drug interactions range from antagonism (left, low *α*) to synergy (right, high *α*). Combining antibiotics with different *α* results in a dose-response surface of more complicated shape (middle). (c) Phase diagram of drug interactions: *LI* score for dose-response surfaces of antibiotic pairs with response parameters *α_A_, α_B_*; white dashed line shows additive interactions (*LI* = 0). The left- and right-hand antibiotic pairs from (b) are shown by a purple triangle and a purple square, respectively. Experimentally determined examples kasugamycin-streptomycin (KSG-STR, antagonistic) and tetracycline-erythromycin (TET-ERM, synergistic) are shown by a white square and circle, respectively. Response parameter values for KSG-STR and TET-ERM are from Ref. [12].

The fact that combinations of antibiotics that bind the ribosome irreversibly yield antagonism can be understood intuitively. If a low concentration of an irreversibly-binding antibiotic is added to a bacterial population in the presence of a high concentration of another such antibiotic, it will mostly bind ribosomes that are already tightly bound by the other antibiotic. Such double-binding events will not lower the growth rate further since the ribosomes were already inactivated by the first antibiotic that bound. Irreversibly-bound ribosomes thus effectively act as a “sponge” that soaks up antibiotics which can then no longer contribute to growth inhibition –a situation that results in antagonism.

What causes the transition from antagonism to synergy as *α* increases? Increasing *α* implies that the binding of the antibiotic becomes more and more reversible (*KD* → ∞), so that the antibiotic-ribosome binding reaction can rapidly equilibrate. In this regime, we can derive an approximate solution that yields a synergistic dose-response surface, supporting the conclusion that qualitatively changing the binding kinetics alters the drug interaction type. If we assume, for simplicity, that λ_0_ ≈ Δ*rκ_t_* = λ_max_, which removes the upregulation of ribosomes since *r*_tot_ ≈ *r*_max_ (or that growth is inhibited close to zero, which also leads to maximal upregulation), and that the time scale of dilution by growth is much greater than that of antibiotic-ribosome binding, we obtain *a* ≈ *a*_ex_*p*_in_/*p*_out_. The system becomes linear and analytically solvable (see Appendix D). The growth rate is then:

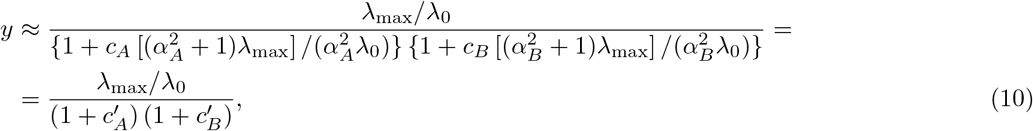

where 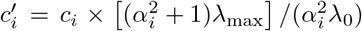. This expression for *y* is simply a product of relative responses, which is equivalent to the definition of Bliss independence. This approximate solution agrees well with the full numerical solution at lower growth rates and for antibiotics with higher *α* (Fig. 4). Equation (10) becomes even simpler in the limit λ_0_ = λ_max_ and *α* → ∞ as these two limits yield a product of two Langmuir-like equations with only relative concentrations *c_A_, c_B_* as arguments, *i.e*., *y* = 1/ [(1 + *c_A_*)(1 + *c_B_*)], which is independent of λ_0_ and *α*.

**FIG. 4.**
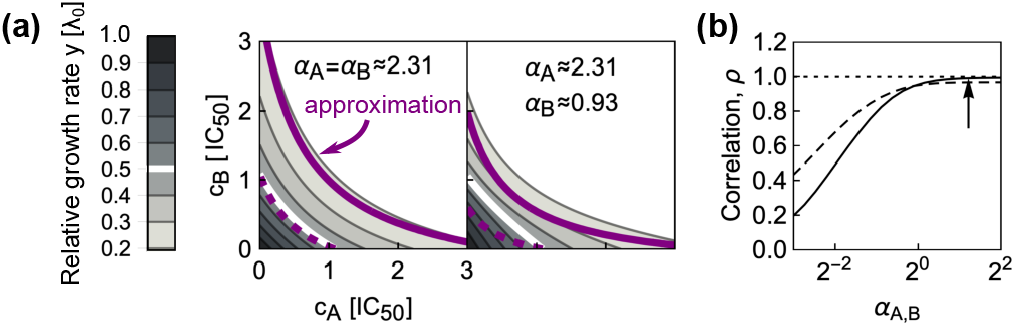
Combining antibiotics with rapidly reversible ribosome binding yields synergistic drug interactions. (a) Comparisons of numerically calculated dose-response surfaces and approximate solution. Purple isoboles (dashed and solid lines correspond to 50% and 20% relative growth rate, respectively) show the approximate solution on top of the dose-response surface calculated from the biophysical model (gray scale). Examples are shown for two pairs of antibiotics with identical (left) or different *α* (right). (b) Pearson correlation *ρ* between approximate and numerically calculated growth rate, evaluated for 121 × 121 equidistant concentration pairs. Solid and dashed line correspond to the cases with identical or different *α*, respectively. The correlation increases for antibiotics with higher *α*. The arrow shows *α* for the example on the left in (a).

### B. Symmetric direct interactions on the ribosome amplify drug interactions

We next asked how more general binding schemes, in which two different antibiotics can directly interact on the ribosome to stabilize or destabilize their binding affect the resulting drug interactions. Two antibiotics do not need to come into direct, physical contact to affect each other’s binding: Allosteric effects (*i.e*., changes in ribosome structure due to antibiotic binding) can produce the same result. In the most plausible scenario, the antibiotics affect each other’s binding in a symmetric way, *i.e*., 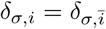 (Fig. 5a). For example, the antibiotics lankamycin and lankacidin (which interact synergistically [19]) are near each other when bound to the ribosome; their binding is stabilized by a direct physical interaction [20, 21]. Yet, it is unclear if the mutual stabilization of binding necessarily leads to synergy. In principle, stabilization of binding could also increase the sequestration of tightly binding antibiotics or “lock” an antibiotic that would rapidly unbind on its own in the bound state, thus potentially promoting prolonged inhibition of the ribosome. In contrast, the two antibiotics may also mutually destabilize their binding to the ribosome. The limiting case of this scenario is competition for the same binding site. To investigate such effects systematically, we computed dose-response surfaces for antibiotics with different response parameters *α* and varying kinetics for the second binding step.

**FIG. 5.**
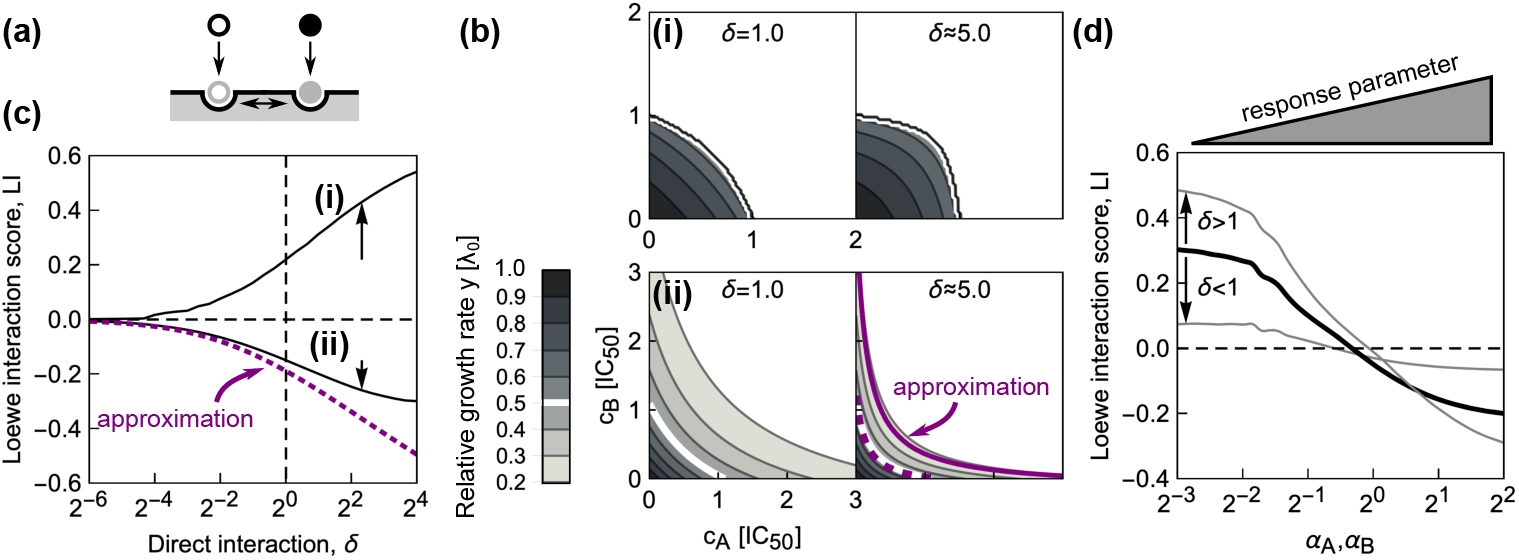
Direct interactions between antibiotics on the ribosome can amplify drug interactions. (a) Schematic of antibiotics symmetrically affecting their binding on the ribosome. (b) Changes in the shape of the dose-response surfaces for pairs of antibiotics with (i) identical *α* = 2^-5^ and (ii) identical *α* = 2^2^, when *δ* is increased from 1 (left) to ≈ 5.0 (right). Purple dashed and solid line in the bottom-right panel show the approximate solution in Eq. (13) for the 50% and 20% isoboles, respectively. (c) Increase in absolute value of the *LI* score as a function of *δ* for pairs of antibiotics with different response parameters. Solid lines (i) and (ii) correspond to the examples in (b); arrows show increase in |*LI*| at *δ* ≈ 5.0. Note, that for both antibiotic combinations, *LI* collapses to 0 for competitive binding, *i.e*., *δ* = 0. The dotted line shows the *LI* score calculated using the approximate solution in Eq. (13). (d) Diagonal cross-section (*α_A_* = *α_B_*) through the phase diagram for different *δ*. Black solid line corresponds to the case of independent binding (*cf*. Fig. 3c); the two gray lines show examples with either *δ* < 1 or *δ* > 1. Irrespective of drug interaction type, the drug interaction is amplified for *δ* > 1 and weakened for *δ* < 1.

We focused on pairs of antibiotics in which both drugs either have low or high *α*, corresponding to steep or shallow dose-response curves, respectively. Numerical solutions for continuously varying *δ*_on,*i*_ = *δ* at fixed *δ*_off,*i*_ = 1 (Fig. 5b,c) showed that a stabilizing interaction (*δ* > 1) enhances the resulting drug interaction. If the drug interaction is antagonistic for *δ* = 1, stabilization amplifies this antagonism; synergistic interactions are amplified analogously (Fig. 5b). If one antibiotic destabilizes the binding of the other, *i.e*., *δ* < 1, a smooth transition to additivity occurs, independent of whether the dose-response curve of the antibiotic pair is steep or shallow (Fig. 5c). This result is further corroborated by fixing *δ* and continuously varying *α* for the combined antibiotics (Fig. 5d). Taken together, these numerical results indicate that direct positive interactions of translation inhibitors on the ribosome (*δ* > 1) essentially amplify the drug interaction that occurs in the absence of such direct interactions, irrespective of drug interaction type.

To corroborate these numerical results, we investigated the limit of reversibly binding antibiotics with rapid binding kinetics at low growth rates as for Eq. (10). In this limit, there is again an analytical solution for the dose-response surface:

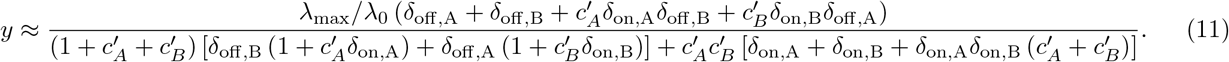

This closed-form expression facilitates the analysis of several limiting cases. For example, if the antibiotics mutually stabilize their binding to the extreme extent that they cannot detach from the double-bound ribosomes anymore (*δ*_off,*i*_ = 0), Eq. (11) returns *y* = 0 indicating strong synergism. In contrast, prohibiting the formation of double-bound ribosomes by setting *δ*_on,*i*_ = 0, yields

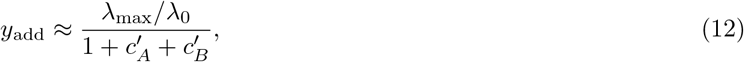

which corresponds to perfect additivity. This corroborates the previous result that competitively binding antibiotics interact additively. For the case *δ*_on,*i*_ = *δ* and *δ*_off,*i*_ = 1 the expression in Eq. (11) simplifies to:

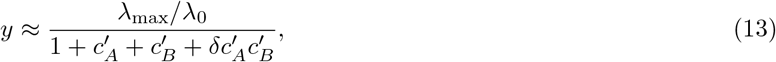

which becomes Eq. (10) if *δ* =1. We can further show that the effect of increasing *δ* on drug interaction strength depends on the concavity of the individual dose-response curves; for response parameters *α* > 2, increasing *δ* amplifies synergy (see Appendix E). Overall, this analysis corroborates the general result that direct stabilizing interactions of reversibly-binding antibiotics on the ribosome amplify synergistic interactions, while destabilizing interactions weaken them up to the point where any drug interaction becomes additive.

### C. Asymmetric direct interactions alter the phase diagram

More generally, direct interactions between the antibiotics on the ribosome could be asymmetric. For example, binding of only one of the antibiotics could trap the ribosome in a conformation that facilitates the binding of the other antibiotic but not *vice versa*. To investigate such effects, we fixed *δ*_off,*i*_ = 1 and varied *δ*_on,*i*_ for antibiotics with different response parameters α (Fig. 6). The resulting difference in kinetic parameters describes an asymmetric direct interaction on the ribosome. We systematically calculated the shape of the dose-response surface for this situation.

**FIG. 6.**
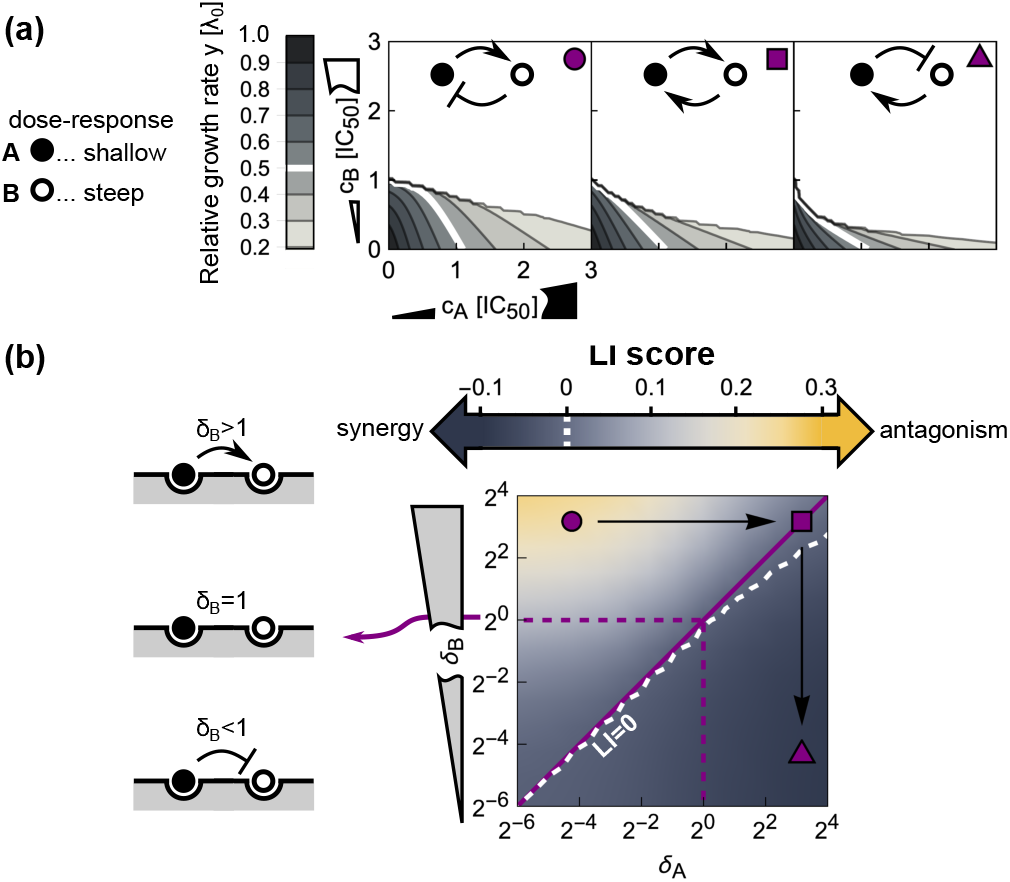
Asymmetric direct interactions reshape the phase diagram of drug interactions. (a) Dose-response surfaces for different instances of asymmetric direct interaction and response parameters; insets on top show schematics of the type of direct interaction and top-right symbols correspond to those in (b). Antibiotics with shallow (*α_A_* = 2^2^) and steep (*α_B_* = 2^-3^) dose-response curves are shown by black and white disks, respectively. Left: Antagonism occurs when an antibiotic with a steep dose-response asymmetrically hinders the binding of another one with a shallow dose-response, which in turn promotes the binding of the former. Middle: Symmetrizing the direct interaction almost completely abolishes antagonism. Right: Inverting the scenario from the left-most panel results in mild synergy. (b) Phase diagram of drug interactions for asymmetric direct interactions between antibiotics with different response parameters. Different response parameters (*α_A_* = 2^2^ and *α_B_* = 2^-3^) profoundly affect the resulting drug interaction: A continuous transition from antagonism to synergy occurs (white dashed line denotes *LI* = 0). Purple symbols show the examples from (a) in the phase diagram.

When antibiotics with identical response parameters *α* are combined, the same trend as for symmetric direct interactions occurs: Increasing *δ*_on_ enhances the drug interaction. For combinations of antibiotics with different doseresponse curve shapes, asymmetric direct interactions on the ribosome result in a different behavior (Fig. 6a,b). If an antibiotic with a steep dose-response curve asymmetrically hinders the binding of an antibiotic with a shallow dose-response, while the binding of the former is stabilized by the latter, antagonism emerges (Fig. 6a). In contrast, synergy occurs if the roles of the antibiotics are inverted (Fig. 6a). The latter can be rationalized by interpreting the direct interaction on the ribosome as a change of the effective binding characteristics of the antibiotics. Specifically, in the case where the steep-response antibiotic promotes the binding of the shallow-response antibiotic, the latter will in turn destabilize the binding of the former–effectively, the steep-response antibiotic will thus behave as if it had a shallower response. As a result, synergy occurs–exactly as expected when two shallow-response antibiotics are combined (Fig. 3c). In the opposite situation, the binding of the shallow-response antibiotic becomes even looser and the binding of the steep-response antibiotic is stabilized. From the phase diagram in Fig. 3c, antagonism is the expected outcome in this case as we combine antibiotics with steep and shallow responses, respectively. Taken together, these results show how complicated direct interactions between antibiotics bound to their target can lead to unexpected emergent drug interactions.

### D. Relation to mechanism-independent models of higher-order drug interactions

The biophysical model described above can predict the pairwise drug interactions that are needed to apply recently proposed mechanism-independent models for higher-order drug interactions [8]. While a detailed analysis of higher-order drug interactions is beyond the scope of this article, it is instructive to demonstrate how the pairwise interactions bridge the gap between responses to individual drugs and higher-order drug combinations. In the framework of Ref. [8], higher-order drug interactions can be predicted using an entropy maximization method, in which the joint drug effects are fully determined by the responses to the individual drugs (*y_i_*) and their pairwise combinations (*y_ij_*):

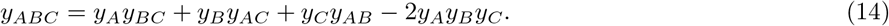

In the limit of reversibly binding antibiotics used in Sec. III A and III B, it is straightforward to analyze the effects of higher-order drug combinations. The approximate results below are based on the assumptions that: (i) the ribosome concentration is already near *r*_max_; (ii) the intracellular concentration of the antibiotic is simply proportional to the external antibiotic concentration; and (iii) the growth rate is directly proportional to the concentration of unblocked ribosomes. Under these assumptions, analytical solutions can be obtained. For example, we can construct a system of differential equations describing the binding of three different antibiotics (*A, B, C*); the steady-state solution of this system is:

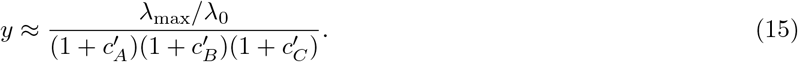

To derive this result, we considered three different kinds of single- and double-bound ribosomes as well as triple-bound ribosomes; for simplicity, all binding steps were considered to be independent of already bound antibiotics. By plugging Eq. (15) into Eq. (14), we note that the mechanism-independent model is consistent with the approximate solution of our model for a combination of three drugs.

Next, we tested if the mechanism-independent formula for three drugs [Eq. (14)] can be reconciled with our model when there is one direct competitive interaction on the ribosome. If the antibiotics *B* and *C* cannot bind to the ribosome simultaneously, the solution of the approximate system is:

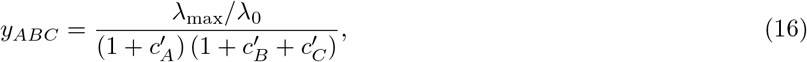

which is again consistent with the mechanism-independent model in Eq. (14). The assumptions described above can be generalized to antibiotics with other modes of action, provided that the binding of the antibiotic to the target is reversible, the growth rate is small compared to the binding rates and proportional to the concentration of the unbound target. These results provide a potential mechanistic explanation for the apparent validity of the mechanism-independent model, at least for combinations of antibiotics binding the same target: The resulting growth rate is invariant of the exact target details as it can be recovered from first-order kinetics.

## IV. EXTENSIONS OF THE MODEL

Other phenomena than those treated so far can shape drug interactions. Below, we discuss two cases in which (A) antibiotics perturb translation in orthogonal ways and (B) the expression of antibiotic resistance genes alters a drug interaction. While certainly not exhaustive, these two cases illustrate relevant extensions of the model.

### A. Effects of antibiotic-induced starvation

Translation inhibitors target the protein synthesis machinery, which is carefully regulated in response to changes in the nutrient environment [22]. Thus, if an antibiotic effectively interferes with cellular state variables that represent the nutrient environment, it should be possible to predict its effect on the action of a translation inhibitor and, in turn, its drug interaction with a translation inhibitor.

Bacterial growth strongly depends on the availability and quality of nutrients. Protein synthesis requires that amino acids are delivered to the translation machinery (ribosomes) by dedicated proteins [elongation factors (EF-Tu)] [23]. The latter bring charged tRNAs (*i.e*., tRNAs with an attached amino acid) to the ribosome (Fig. 7b). tRNAs are charged (*i.e*., amino acids are attached to them) by tRNA synthetases. Usually, the supply and demand of amino acids can be considered to be nearly optimally regulated [22] (Fig. 7a). However, under starvation, a mismatch between the supply and demand of amino acids occurs [24]. Bacteria respond to amino acid starvation by triggering the stringent response. This starvation response is primarily controlled by the alarmone ppGpp (guanosine tetraphosphate) which down-regulates the expression of the translation machinery (Fig. 7a) [25]. Amino acid starvation is reflected in reduced tRNA charging and usually occurs when the nutrient environment becomes poor. However, amino acid starvation can also be caused by a *starvation-mimicking antibiotic* (SMA) that blocks tRNA synthetases (Fig. 7b) [26, 27].

**FIG. 7.**
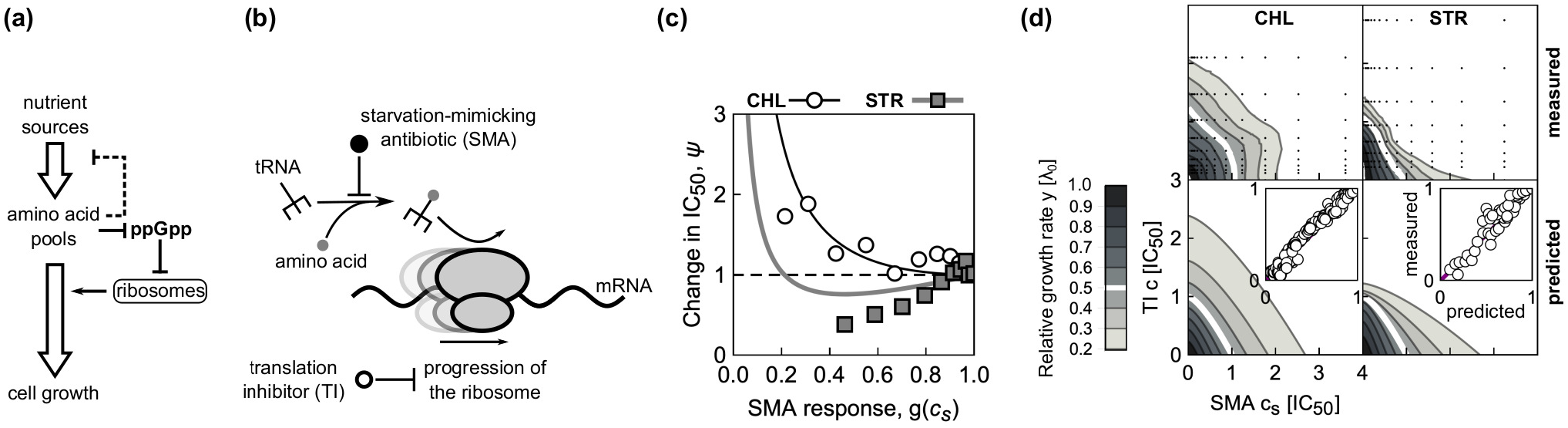
Effects of a starvation-mimicking antibiotic on the efficacy of translation inhibitors. (a) Schematic: Simplified regulation of translation coordination. Nutrients are transported into the cell, where they serve as a source of amino acids. These amino acids are required for tRNA charging. Oversupply of amino acids leads to down-regulation of the nutrient transport and processing machinery, and depletion of the intracellular signaling molecule ppGpp (guanosine tetraphosphate). This in turn de-represses the expression of the translation machinery, which increases the overall translation capacity, leading to faster growth. In contrast, if amino acids are in short supply, the translation machinery is down-regulated. (b) Translation inhibitors (TI) inhibit progression of the ribosome, while a starvation-mimicking antibiotic (SMA) perturbs the amino acid supply. The ribosome progresses along the mRNA (black wavy line), if charged tRNAs (black fork with gray circle) deliver amino acids (gray circles) at a sufficient rate to support the rapid synthesis. A starvation-mimicking antibiotic inhibits tRNA charging and thus mimics amino acid depletion, a hallmark of starvation. (c) Dependence of relative change in IC_50_ on SMA inhibition (*ψ* = IC_50_/IC_50,*F*_). Example solutions of Eq. (17) were calculated for chloramphenicol (CHL; black line with *α_F_* = 1.04, white circles show experimental data) and streptomycin (STR; gray line with *α_F_* = 0.46, gray squares show experimental data). In the experiments, mupirocin (MUP) was used as SMA. The horizontal dashed line indicates no change with respect to *g*(*c_s_*). Values for *α* are from Ref. [12]. (d) Measured (top) and predicted (bottom) dose-response surfaces for CHL-MUP (left) and STR-MUP (right). Insets show scatter-plots of predicted and measured non-zero growth rates.

We can capture the effect of an SMA in our model and thus make predictions for the drug interactions between an SMA and translation inhibitors. To this end, we assume that the growth rate in the absence of drug λ_0_, which characterizes the quality of the nutrient environment in Eq. (4), depends on the concentration *c_s_* of the SMA only. Under this assumption, the growth rate in the simultaneous presence of an SMA and a translation inhibitor can be derived directly from the previous results for a single antibiotic [Eq. (5)]. In the absence of translation inhibitor, the growth rate is given by the dose-response function of the SMA *g*(*c_s_*). Since IC_50_ *α*: (*α*^2^ + 1) /*α* and *α* = *α_F_*/*g*(*c_s_*), the relative change in IC_50_ at constant SMA concentration becomes

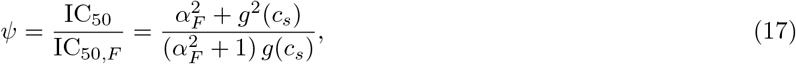

where *α_F_* and IC_50,*F*_ are *α* and IC_50_ in the absence of the SMA, respectively. It follows that *ψ* increases monotonically with SMA inhibition if *α_F_* > 1; this condition is obtained by solving *∂_g_ψ* = 0 for *g*(*c_s_*) ≤ 1. If *α_F_* ≤ 1, then the minimal *ψ* is reached at *g*(*c_s_*) = *α_F_*. The dose-response curve for a single antibiotic is given by *y* = *f*(*α, c*) [a solution of Eq. (5)]. Since we know how IC_50_ [Eq. (17)] and *α* change as a function of *g*(*c_s_*), we can evaluate the entire dose-response surface:

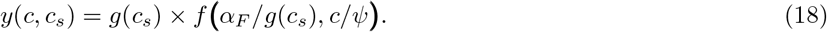

Equation (18) deviates from a simple multiplicative expectation since the SMA affects the parameters of the doseresponse curve of the translation inhibitor as well. This result illustrates how deviations from Bliss independence occur when the combined drugs mutually affect their dose-response characteristics. Hence, this generalization of our model makes non-trivial quantitative predictions for drug interactions that occur between an SMA and translation inhibitors.

A specific example of an SMA is the antibiotic mupirocin (MUP), which reversibly binds to isoleucin tRNA synthetase and prevents tRNA charging [28]. MUP, which is used against clinically problematic methicillin-resistant *Staphylococcus aureus* (MRSA) infections [29], induces the stringent response [26, 27] and can thus be used to test our theoretical prediction. To this end, we measured the change of the IC_50_ at different levels of growth inhibition caused by MUP *g*(*c_s_*) for two translation inhibitors: chloramphenicol (CHL) and streptomycin (STR; see Appendix F). CHL and STR have extremely different response parameters *α* [12], which leads to different dependencies of *ψ* on SMA inhibition (Fig. 7c), an effect that is closely related to the results for different nutrient environments in Ref. [10]. Similarly, we measured the complete dose-response surfaces for both drug pairs. The theoretically predicted doseresponse surfaces qualitatively agree with the experimentally observed ones (Fig. 7d). In Appendix G we discuss further examples of pairwise antibiotic combinations in which a translation inhibitor is combined with a drug that alters the growth law parameters. Together, these results illustrate how our theoretical model can be extended to predict the effects of drug combinations beyond antibiotics that directly target the ribosome.

### B. Effect of constitutively expressed resistance genes

Our results show that the steepness of the dose-response curve and the coupling between growth laws and antibiotic response play a key role in determining drug interactions. Recent work showed that dose-response curve steepness can change if genes that convey antibiotic resistance are present [17]. Thus, we investigated how the presence of such resistance genes affects the resulting drug interaction.

Bacterial resistance genes often code for dedicated enzymes that degrade the antibiotic or pump it out of the cell. Resistance genes can be constitutively expressed, *i.e*., they lack specific regulation and their expression depends only on the state of the gene expression machinery. The expression of such constitutively expressed resistance genes (CERGs) under translation inhibition is quantitatively predicted by a theory based on bacterial growth laws [9, 17]: Expression *q* decreases linearly with decreasing growth rate as *q* = *q*_0_λ/λ_0_, where *q*_0_ is the expression of the gene in the absence of the drug. An experimentally verified mathematical model that is based on this dependence predicts growth bistability (*i.e*., coexistence of growing and non-or slowly-growing cells) in bacterial populations that constitutively express resistance genes [17]. In this model, the flux of antibiotic removal due to the resistance enzyme is described by

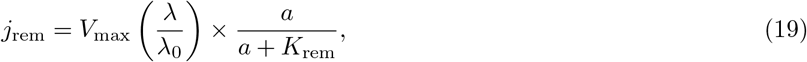

where *K*_rem_ is a Michaelis-Menten dissociation constant and *V*_max_ is the maximal antibiotic removal rate, which bundles the maximal enzyme abundance and catalytic rate per enzyme.

Due to the linear relation between growth rate and the expression of the resistance gene, the rate of antibiotic removal decreases with decreasing growth rate under translation inhibition. This constitutes a positive feedback loop that leads to growth bistability (Fig. 8a,b), which is reflected in a steep dose-response curve of bacterial batch cultures [17]. However, note that for very high values of *k*_rem_ ≫ *a*, Eq. (19) becomes linear and the steepness of the dose-response curve decreases, rendering the otherwise bistable system monostable (see Appendix A 2).

**FIG. 8.**
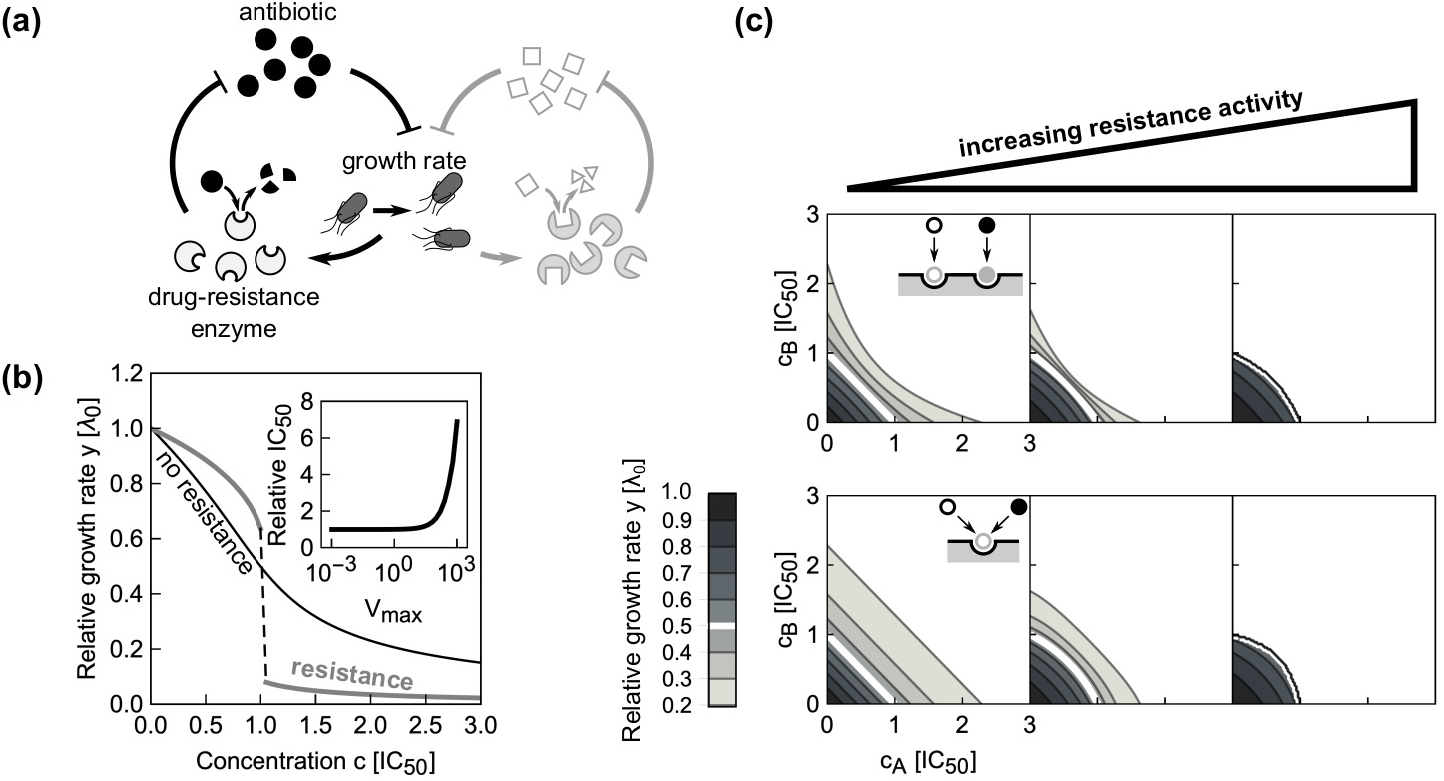
Effects of constitutive resistance genes on the shape of dose-response curves and surfaces. (a) Schematic of positive feedback loop for unregulated antibiotic resistance gene. A drug-resistance enzyme degrades the antibiotic, thus reducing growth inhibition and boosting its own expression. However, if the antibiotic concentration exceeds the capacity of removal by the enzyme, growth rate starts to drop and so does the expression of the resistance enzyme, amplifying the growth rate drop. Lightly drawn part (right) illustrates how two antibiotics can get coupled via the growth-rate dependent loop. (b) Examples of dose-response curves in the presence or in the absence of a constitutively expressed resistance gene (CERG). Black line shows dose-response curve for *α* = 1. When a CERG is added (*V*_max_ = 1000 *μ*M h^−1^, *k*_rem_ = 0.1 *μ*M), the dose-response curve becomes steeper and exhibits an abrupt drop. Inset shows the increase in antibiotic concentration required to halve the growth rate relative to the no-CERG case as a function of *V*_max_. (c) Dose-response surface for independently (top) and competitively (bottom) binding antibiotics with *α* =1; resistance activity *V*_max_ (assumed to be identical for both antibiotics) increases from left to right: 0, 100 and 950 *μ*M h^−1^. Concentration axes were rescaled with respect to the increased IC_50_. Note the qualitative change in dose-response surface shape.

By extending this scenario to a pair of antibiotics, we can directly test how the presence of resistance genes affects drug interactions. In the most relevant case, there are two CERGs each of which specifically provides resistance to one of the antibiotics. For simplicity, we assume that there is no cross-resistance, *i.e*., each enzyme specifically degrades only one of the drugs (Fig. 8a). We found that the synergistic interaction between two independently binding antibiotics with shallow dose-response curves turns slightly antagonistic due to the presence of resistance genes (Fig. 8c, top). For competitively binding antibiotics, this effect becomes more pronounced (Fig. 8c, bottom). In brief, our model predicts drastic qualitative changes in drug interaction type when resistance genes are present.

To test this prediction, we constructed a bacterial strain (see Appendix F) that carries two constitutively expressed resistance genes. We chose TetA [a tetracycline (TET) efflux pump] and CAT [an enzyme that degrades chloramphenicol (CHL)], which were previously characterized in the context of bacterial growth laws (Fig. 9a; [17]). Furthermore, the interaction between CHL and TET is additive. Our model predicts this interaction to change into antagonism when CERGs are present. Consistent with previous results [17], the steepness of the dose-response curve increased upon inclusion of each CERG (Fig. 9b). We measured the dose-response surface of the sensitive and the double-resistant strain: Notably, the resistant strain showed a clear antagonistic drug interaction, while this interaction was additive in the strain without CERGs (Fig. 9c). This change to antagonism qualitatively agrees with the theoretical prediction (Fig. 8c). This example shows how resistance genes can drastically alter drug interactions–a phenomenon caused by a non-trivial interplay of gene-expression and cell physiology predicted by our biophysical model.

**FIG. 9.**
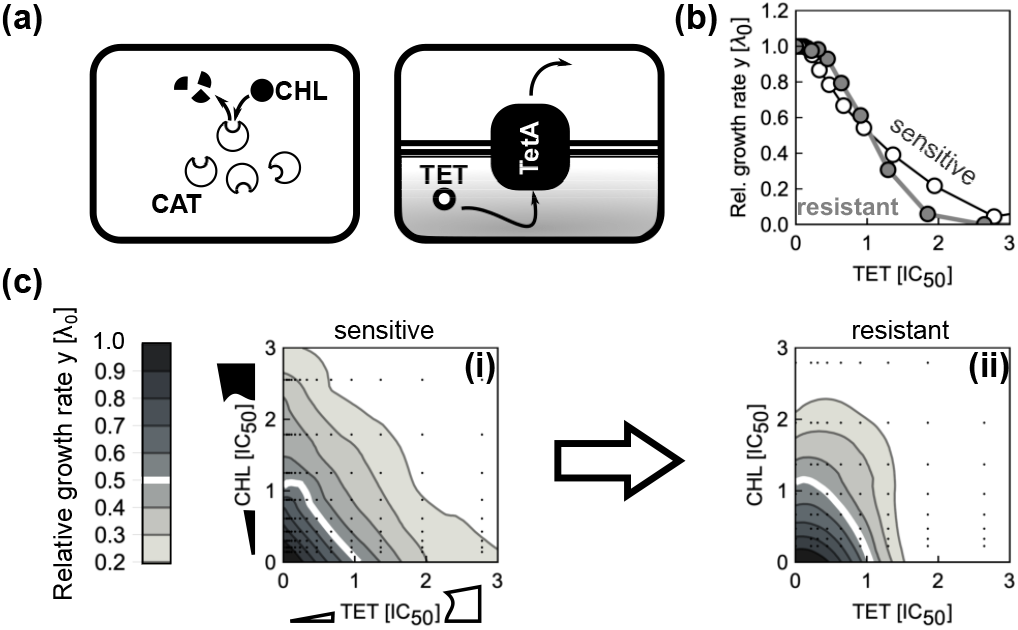
Constitutively expressed resistance genes alter a drug interaction as predicted by theory. (a) Schematic showing two common resistance mechanisms: Resistance can result from degradation of the drug [left: chloramphenicol acetyltransferase (CAT) degrades chloramphenicol (CHL)] or from drug efflux [right: an antibiotic efflux pump (TetA) removes tetracycline (TET) from the cell]. (b) Change in dose-response curve shape due to a constitutively expressed resistance gene. CHL doseresponse curves of sensitive (white circles) and resistant strain (gray circles). (c) Measured CHL-TET dose-response surfaces for (i) sensitive and (ii) resistant strain. Concentrations were normalized to the IC_50_ of respective strains. The strain with CERGs is 50.5 and 91.5 times more resistant to TET and CHL, respectively, as measured by increase of IC_50_. Drug interaction changes from additive to antagonistic as suggested by theory (Fig. 8c).

## V. DISCUSSION

We constructed a minimal biophysical model of antibiotic interactions that takes into account the laws of bacterial cell physiology. Most parameters in our model are constrained by established results or by the dose-response curves of the individual antibiotics that are combined (Fig. 3). Our approach offers a scalable theoretical framework for predicting drug interactions: The number of parameters required for the independent binding model scales linearly with the number of antibiotics. This framework is readily generalized to combinations of more than two antibiotics. Ribosomal growth laws [9] were essential for building this predictive framework, highlighting the importance of quantitative phenomenological descriptions of physiological responses to drugs and other perturbations (Fig. 2). The discovery of similar quantitative relations between physiological parameters and growth rate for other classes of antibiotics and other types of cells would greatly facilitate more general predictions of drug interactions.

Our work highlights the advantages of a physiologically relevant “null model,” which captures all effects that are generally relevant for ribosome-binding antibiotics without trying to describe any molecular details of specific antibiotics (Fig. 3). While general multiplicative (Bliss) or additive (Loewe) expectations are simple to construct, our work demonstrates that their utility as a reference has clear limitations. Specifically, our model shows that both are expected to be valid only in certain limits (Figs. 4 and 5). Moreover, these standard null models do not capture known effects of antibiotic binding and growth physiology, which suffice to produce strong deviations from the standard null models. Our biophysical model captures these effects and thus offers an improved expectation for drug interactions. Generalizing this model to three drugs demonstrated that mechanism-independent predictions of higher-order interactions [8] are consistent with simplified first-order kinetics. In summary, our model serves as a bridge between mechanism-independent general predictions of drug interactions and elusive quantitative descriptions of detailed molecular mechanisms that capture the idiosyncrasies of each drug.

We showed that direct physical (or allosteric) interactions of antibiotics on their target do not necessarily lead to synergy (Figs. 5 and 10). Synergy only occurs if the dose-response curves of the individual drugs are sufficiently shallow. While this insight is not easily applied in the design of drug combinations, the identification of cooperatively binding drug pairs still has considerable potential. Our results highlight that altering the steepness of individual drug dose-response curves may offer under-appreciated opportunities for drug design.

**FIG. 10.**
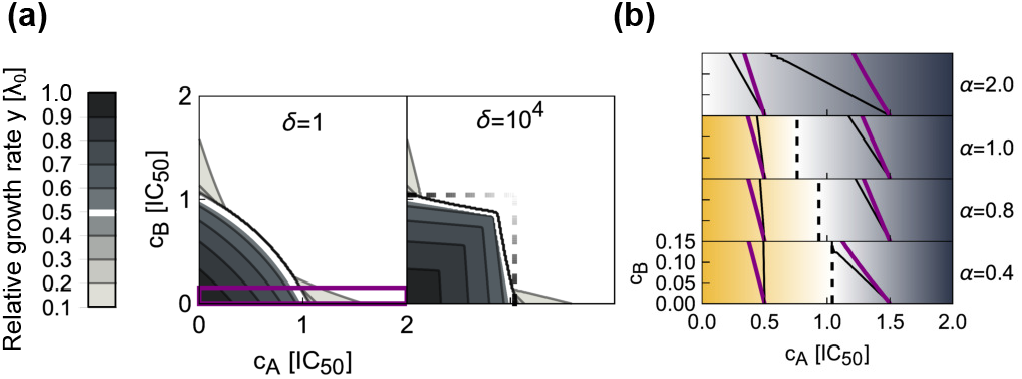
Dose-response surface can locally have both antagonistic and synergistic isoboles, depending on the concavity of the individual dose-response curves. (a) Examples of dose-response surfaces with *α* = 0.4 and with either independent (left, *δ* = 1) or strongly cooperative binding (right, *δ* = 10^4^). Purple box highlights the area showcased in (b). Dashed black lines denote an approximate area in which increasing *δ* increases antagonism. (b) A detail [highlighted rectangle in (a)] of dose-response surfaces for different values of *α*. For *α* < 2 black dashed line denotes the boundary below which the increasing cooperativity *δ* increases antagonism (shades of yellow); above it synergistic character is enhanced (blue). Two isoboles are showcased for each example: purple and black lines correspond to *δ* = 1 and *δ* = 10^4^, respectively. Note, that below *c*_inf_ isoboles in the case of strong cooperativity become steeper than the independent ones; above *c*_inf_ isoboles become shallower, which is indicative of synergy.

The predictions of our model are directly testable in experiments (Section IV and Ref. [12]). Perhaps the most striking experimental validation of our model is the change in drug interaction type due to the presence of antibiotic resistance genes (Fig. 8c, Fig. 9c). This observation is notable since previous work concluded that most mutations and mechanisms that provide resistance to individual drugs only rescale the effective antibiotic concentrations while preserving the shape of the dose-response curves and surfaces [30–33]. In contrast, our results show that specific resistance genes for two antibiotics targeting the ribosome inevitably alter the drug interaction, even in the absence of more complicated mechanisms.

Discrepancies between experimental results and model predictions can expose cases in which more complicated mechanisms cause the observed drug interaction [12]. A limitation of our model is that it considers fully assembled translating ribosomes as sole targets of the antibiotics, without taking the exact stage of the translation cycle into account. In principle, a model that describes ribosome assembly and more details of the translation cycle and the transitions between its different steps could provide a more detailed mechanistic picture. However, since we currently do not know the *in vivo* parameter values that characterize the translation cycle, such a model would not be predictive, but would rather rely on extensive fitting of free parameters to limited experimental data. Instead, the underlying mechanisms of drug interactions that cannot be captured by the minimal biophysical model presented here, and in particular suppression, can be elucidated by targeted phenomenological approaches [12].

## ACKNOWLEDGMENTS

This work was supported in part by Tum stipend of Knafelj foundation (to B.K.), Austrian Science Fund (FWF) standalone grants P 27201-B22 (to T.B.) and P 28844 (to G.T.), HFSP program Grant RGP0042/2013 (to T.B.), German Research Foundation (DFG) individual grant BO 3502/2-1 (to T.B.), and German Research Foundation (DFG) Collaborative Research Centre (SFB) 1310 (to T.B.). B.K. is thankful to C. Guet for additional guidance and generous support which rendered this work possible. B.K. thanks all members of Guet group for many helpful discussions and sharing of laboratory resources. Authors thank M. Hennessey-Wesen and M. Zagorski for reading the manuscript and constructive comments.

## Appendix A: Parameter reduction and the bifurcation point

In the following we derive expressions that constrain the parameter space in which the system is bistable. We start by rewriting the steady-state solution with new variables, which will facilitate reduction of parameters. We recast Eq. (4) so as to express 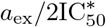 as function of growth rate; this is possible since 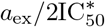 occurs linearly in the equation. At this stage we introduce new variables and rewrite parameters in a dimensionless form:

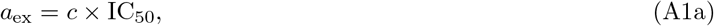

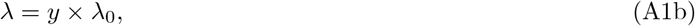

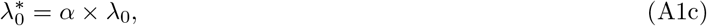

where IC_50_ is concentration needed to halve the growth rate and λ_0_ is a drug-free growth rate. These definitions lead to:

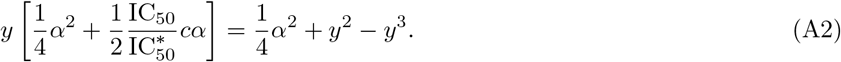

To remove the ratio 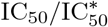 from the equation, we use the expression from Ref. [10] that states

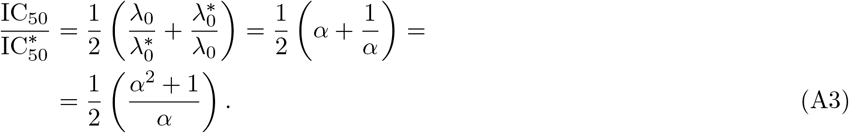

This relation is obtained by plugging *y* = 1/2 and *c* = 1 into Eq. (A2). Finally, we recast Eq. (A2) into

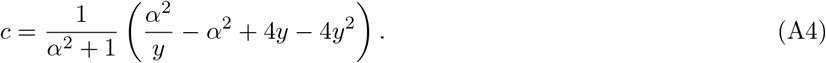

This equation expresses the relative concentration *c* as a function of relative growth rate *y* in which *α* is the sole parameter. Fitting an implicit function can be challenging; we can estimate α from equating the implicitly calculated derivative of Eq. (A4) to the derivative of the Hill function with steepness parameter *n*, when both derivatives are evaluated at *c* = 1. An estimate for the response parameter is then 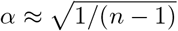.

### 1. Bifurcation point

To determine the critical value of the parameter *α* below which bistable concentration region exists, we consider the stationary points of Eq. (A4). The derivative reads:

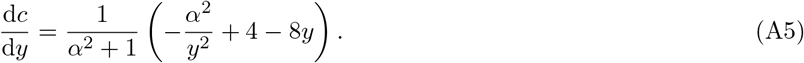

Considering the equation

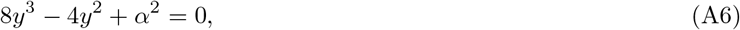

we can determine values of *y* where the derivative of Eq. (A5) is 0. Since the equation is of third order, we expect either three real roots or one real together with a conjugated pair of complex roots. The solutions [found using Mathematica (Wolfram Research, 11.3) function `Solve`] are:

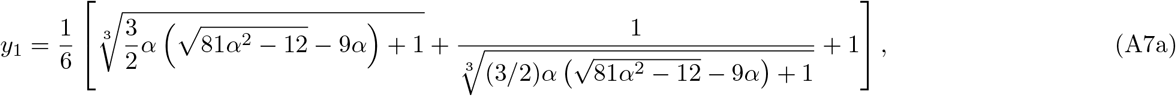

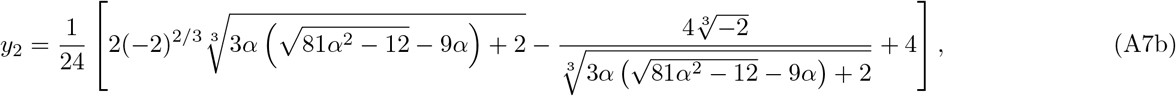

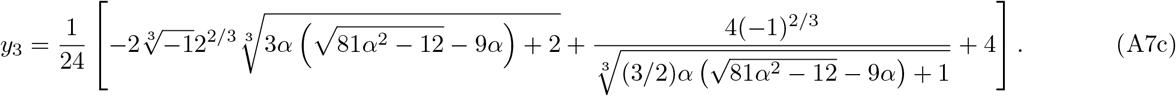

We note that whether all roots are real is determined by the sign of 27*α*^2^ – 4; this leads to a critical value of 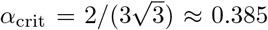–the system has a bistable region when *α* < *α*_crit_. To obtain the span of concentrations at which two bistable growth rates exist (Fig. 2d), we plug Eqs. (A7a) and (A7c) into Eq. (A4).

### 2. Linearized case of constitutively expressed resistance gene

As noted in the main text (Sec. IVB), for high values of *K*_rem_ the Michaelis-Menten equation linearizes. In this case we can reuse the expression for growth-dependent efflux from Ref. [10]. Using the parameter reduction from above, the generic solution is

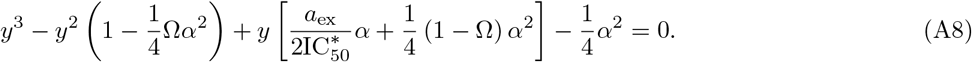

where Ω = *V*_max_/(*p*_out_*K*_rem_). This expression can be rewritten into reduced form as

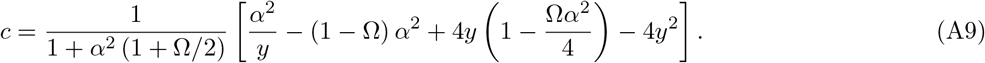

Here, we used the relation that follows directly from Eq. (A8); 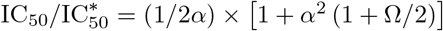. By solving d*c*/d*y* = 0 we obtain a condition −64*α*^2^ + 432*α*^4^ + 48*α*^4^Ω − 12*α*^6^Ω^2^ + *α*^8^Ω^3^ ≤ 0 for bistability to exist; heuristically, we note that the leading term in Ω is of the third power and positive and will for high values of Ω make the expression necessarily positive and will thus make the system monostable. This illustrates the importance of non-linearity in the expression describing antibiotic removal [Eq. (19)]; the presence of positive feedback alone is not sufficient for the occurrence of bistability. In the limit Ω → ∞, Eq. (A9) becomes simply *y* = 1 − *c*/2 for *c* ≤ 2 and *y* = 0 otherwise.

## Appendix B: Calculation of dose-response surface for additive interaction and Loewe interaction score

An additive interaction is characterized by linear isoboles. In this section we briefly sketch the derivation. Let *c_A_* and *c_B_* be relative concentrations (measured in IC_50_ of respective antibiotics) of antibiotics *A* and *B*, respectively. Individual dose-response curves are given by *f_A_* and *f_B_*. To construct the additive surface, the growth rate in any point (*c_A_, c_B_*) has to be calculated solely from known responses to individual drugs. As isoboles are linear, there is an isobole that passes through this point but terminates in some unique (*c*_*A*,0_, 0) and (0, *c*_*B*,0_), from which it follows that *c_B_* = *c*_*B*,0_(1 − *c_A_*/*c*_*A*,0_). Since 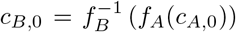, the solution of 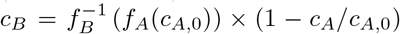 for *c*_*A*,0_ is sought, which depends on *c_A_* and *c_B_*. If such solution is termed 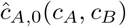 then the whole dose-response surface is given as 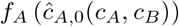. Integration of Eq. (9) is performed over an area where growth rate is above a chosen threshold. Throughout the article this threshold is set to *y* = 0.2. Positive or negative values of *LI* suggest antagonistic or synergistic interaction, respectively.

## Appendix C: Numerical solutions

We evaluated the steady-state solution of the system Eqs. (6) by forward time integration of the differential equations. We used λ_0_ = 2 h^−1^, which is the drug-free growth rate in rich lysogeny broth medium at 37°C [12]. We rescaled the time by defining *t* = *Tτ*, where *T* = (*α_A_* + *α_B_*)^−1^ × (1/λ_0_) and *τ* is a real number. Similarly, we rescaled concentrations by defining a reference concentration as 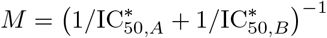; concentrations of all species were rescaled by *M*. To automatically scale the antibiotic concentrations axes in units of IC_50_, we set 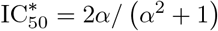. To mimic the situation in which exponentially growing unperturbed bacterial culture is exposed to antibiotic, we set the initial condition to *y* = 1; the rest of the species were set to 0. We integrated the rescaled differential equations forward in time until *τ* = 9 × 10^3^ by using Mathematica function `NDSolve`. In computation, *K_D_* and *k*_on_ values were set for both antibiotics to 0.1 *μ*M and 100 *μ*M^−1^h^−1^, respectively. The results are largely invariant of the choice of these numeric constants as long *k*_on_ ≫ *κ_t_* and *K_D_* is roughly between 0.01 – 1 *μ*M.

## Appendix D: Analytical solution in the limit of strong inhibition and reversible binding

The system of ODEs [Eqs. (6a-d)] can be linearized for near-zero growth rates (*i.e*., 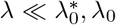) which is the case when the external concentrations of antibiotics are high enough (*a*_ex,*i*_ ≫ IC_50,*i*_). In the latter case one can postulate two constraints: (i) Ribosome synthesis is up-regulated to the theoretical maximum, *i.e*., *r*_tot_ = *r*_max_ and (ii) internal concentration of the *i*-th antibiotic is 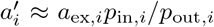. These constraints eliminate the dynamical equations for internal concentrations of antibiotics [Eq. (6a)]. The fully specified and expanded system of equations reads

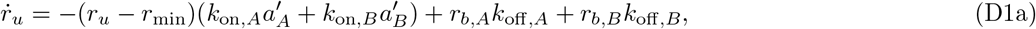

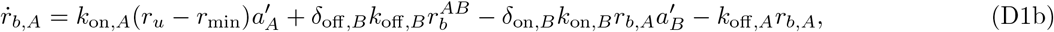

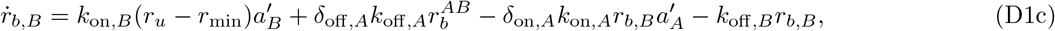

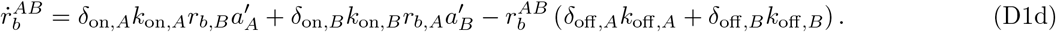

The system is linear and thus the steady-state solution exists in a closed and unique form. We initally derive the stationary solution for independent binding, *i.e*., *δ_σ,i_* = 1:

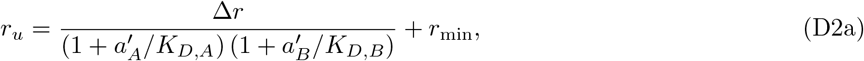

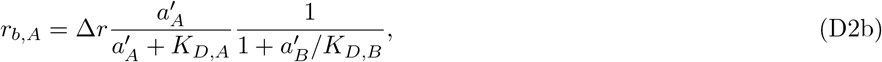

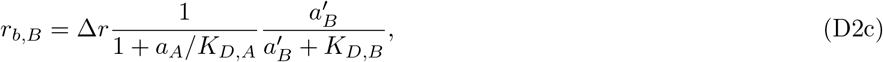

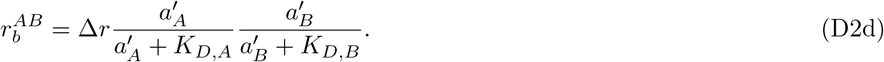

Each product term can be rewritten as a function of 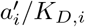, which can be further transformed by noting that 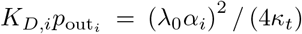 and 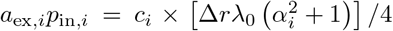. Together, these expressions allow rewriting Eq. (D2a) using variables 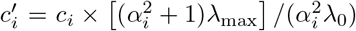, which finally leads to the Eq. (10). To obtain Eq. (11) the general solution of Eqs. (D1) is evaluated; the latter is further simplified by assuming that *k*_off,*A*_ ≈ *k*_off,*B*_, which approximately holds for reversibly binding antibiotics.

## Appendix E: Effect of dose-response curve concavity on the shape of isoboles

For intermediate values of *α* ~ 1 and increasing *δ*, we observed that the isoboles of the dose-response surface at lower drug concentrations indicate strong antagonism, whereas at higher concentrations, they indicate synergism (Fig. 10a). Is it possible to determine a concentration value above which increasing *δ* will cease to increase antagonism but rather increase synergism? Intuitively, the interaction-characteristic shape of isoboles is determined by the sign of the mean curvature, the latter being defined as

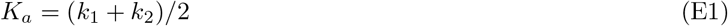

where *k*_1_ and *k*_2_ are principal curvatures. Full dependency of *K_a_* is not accessible as an analytical expression for *y*(*c_A_, c_B_*) is not known; however in the limit *δ* → ∞ the antagonistic isoboles become nearly perpendicular to the axes, thus rendering all derivatives along perpendicular directions equal to zero. Hence, the expression for *K_a_* is simplified into:

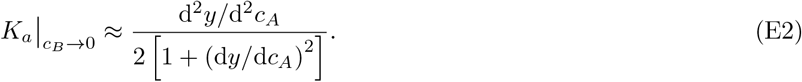

As the denominator is always positive the sign of *K_a_* depends only on the sign of the second derivative along the axis.

To find this point, we need to determine the inflection point of the relative growth rate *y* along either of the axes. Here, we need to solve d^2^*y*/d^2^*c* = 0 for *α*; since *y* is given only implicitly in Eq. (5), the second implicit derivative is calculated. This leads to inflection point at relative growth rate 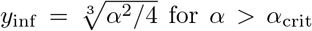 for *α* > *α*_crit_, at which second derivative vanishes. Using Eq. (5) the inflection concentration is evaluated as:

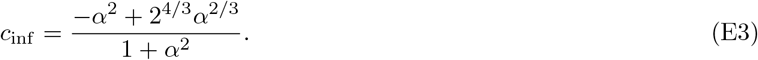

From the expression for *y*_inf_ we observe that we have a relevant solution only for *α* < 2; this leads to the first conclusion–the intermediate regime of surfaces having both antagonistic and synergistic contours exists only up to *α* = 2. This also implies that for every response parameter α, given a high enough concentration of an antibiotic, the isoboles will become synergistic. However, at low growth rates at high concentrations the approximation [Eq. (13)] becomes relevant (Fig. 10b). Numerically computed dose-response surfaces for *δ* = 1 and *δ* = 10^4^ illustrate that indeed above the *c*_inf_ character of the isoboles is different and that the transition happens due to vanishing second derivative.

## Appendix F: Experimental considerations

We verified the results on antibiotic induced starvation and constitutive expression of resistance genes (Secs. IV A and IV B, respectively) by measuring the growth rate in two-dimensional gradients as described in Ref. [12]. Briefly, concentration gradients were constructed in the wells of microtiter plates; multiple such plates were cycled through the plate reader, which measured the luminescence as the proxy of the cell density. From the time traces of luminescence the growth rate of the bacterial culture in the individual well was determined by identifying an exponential region and fitting the line to the log-transformed luminescence values–precise methods and experimental setup description is in Ref. [12]. Bacterial growth medium used in all experiments was lysogeny broth (LB). Antibiotics solutions were prepared from powder stocks and dissolved in ethanol (CHL, MUP), water (STR) or directly in LB [TET and CHL (for CERG experiment)] and filter-sterilized. Growth rates were normalized with respect to the antibiotic-free growth rate λ_0_ to obtain *y* = λ/λ_0_; concentrations were normalized with respect to IC_50_, which was determined from fitting a Hill function 1/ [1 + (*a*_ex,*i*_/IC_50,*i*_)^*n*^]. To construct the double-resistant strain we followed the methods described in Refs. [12] and [17] with appropriate modifications. We have cloned cat gene into a plasmid with pKD13 background (as per Ref. [12]), in which resistance gene was driven by a synthetic promoter P_L1acO-1_ [34]. Promoter was unregulated (constitutive) as the background strain HG105 (MG1655 Δ*lacIZYA*) [35] is devoid of the entire *lac* operon, including the lac-repressor. We amplified the *tetA* and *cat* from the strain MS004A [36] and plasmid pZA32 [34], respectively. We were unable to clone *tetA* into a plasmid; we therefore assembled the kanamycin cassette, P_LlacO-1_ promoter, and *tetA* gene with a *rrnB* terminator *in vitro* using HiFi Assembly Mix (NEB), PCR-amplified the fragment, and integrated it into the *intS* locus. CAT-coding gene was integrated into *galK* locus. Both CERGs were integrated into the chromosome by lambda-red recombineering, selected on kanamycin, which was followed by the resolution of the kanamycin cassette by FLP-resolvase [12, 17, 37–39]. We verified modified chromosomal segments with PCR and Sanger sequencing.

## Appendix G: Combinations of translation inhibitors with drugs that alter growth law parameters

We elaborated in the main text on a specific example of a starvation mimicking antibiotic, which invokes a response that is similar to shifting bacteria to poorer media (*i.e*., change in λ_0_). As growth laws capture this change, a response could be analytically predicted and was experimentally tested. Below we show two more hypothetical examples of antibiotic combinations in which translation inhibitor is combined with a drug that alters growth law parameters. We consider the case in which this hypothetical drug alters either *r*_min_ alone or in concert with *r*_max_; in either case a shift in growth laws happens, which affects both dynamic range Δ*r* as well as the apparent response parameter *α*. Here, we note that there is some experimental data illustrating that such shift could be achieved by antibiotics inhibiting *transcription, e.g*., rifampicin (Fig. 11a and supplementary information of Ref. [9]). However, the combined effect of transcription and translation inhibition of translation machinery has not be experimentally assessed.

**FIG. 11.**
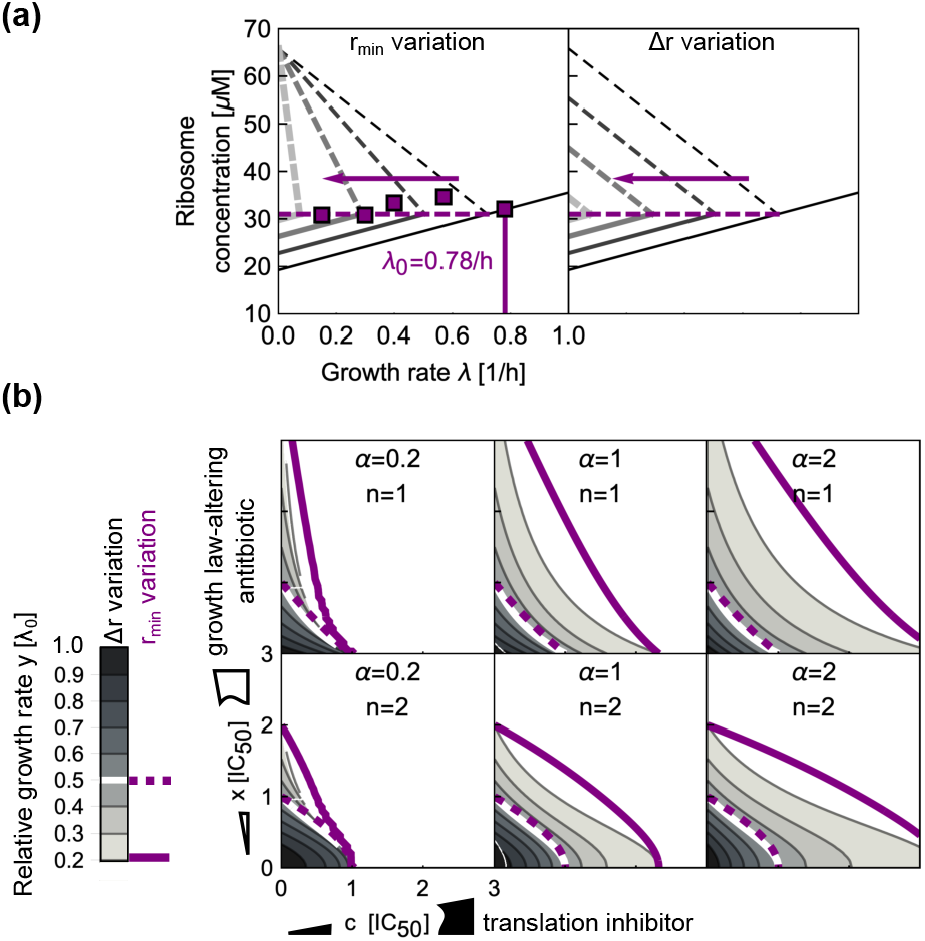
Impact of the variation in *r*_min_ and Δ*r* on drug interactions. (a) Changes in growth laws as a function of variation in *r*_min_ (left) and Δ*r* (right). Purple arrow denotes the effect of growth law-varying drug, which alter the “origin” of the translation inhibition growth law line (dashed gray lines). Slope of dashed lines is altered when *r*_min_ is varied; when *r*_max_ is varied simultaneously with *r*_min_ the slope is invariant of the “origin.” Purple squares is data describing the ribosome abundance in response to rifampicin from Ref. [9]; we converted reported RNA/protein measurements to ribosome concentrations by fixing the drug-free data point to the ribosome concentration *r_u_* ≈ 32.0 *μ*M as determined by the Eq. (1) at λ_0_ = 0.78 h^−1^. Note, that the data points lie approximately on the horizontal dashed line. (b) Examples of dose-response surfaces for different translation inhibitors (concentration c; different response parameters *α*) and different dose-response curves fo*r* Δ*r*- and *r*_min_varying antibiotic (concentration *x*; Hill-function with different steepness parameter *n*). For easier comparison we overlaid the dose-response surfaces; the purple contours correspond to Δ*r*-varying case.

If we consider an antibiotic that affects the *r*_min_, yet it does not affect the *κ_t_* (Fig. 11a), then the following modification to the growth law arises:

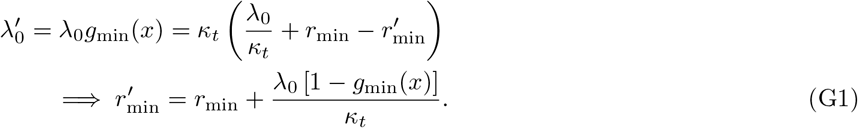

Here, we denoted by *g*_min_(*x*) a dose-response function of a *r*_min_-varying antibiotic and *x* its concentration; 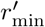 denotes an apparent new *r*_min_ as a function of *g*_min_(*x*). Thus, the reduced dynamic range reads 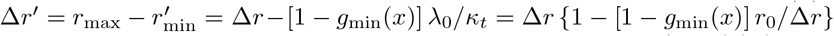, where we defined *r*_0_ = λ_0_/*κ_t_*. Taken together, these relations rescale the response parameter *α* = *α_F_*/*g*_min_(*x*) (where *α_F_* is the response parameter for *x* = 0), while the apparent concentration becomes *c′* = *c*/*ψ*_min_, where

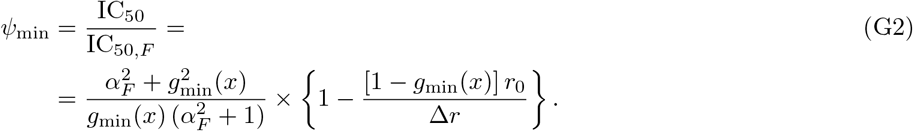

If we consider that *g*_min_(*x*) = 1/ (1 + *x^n^*), where *x* is the concentration of *r*_min_-varying antibiotic measured in units of IC_50_, we can calculate the whole dose-response surface as

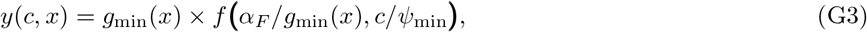

where *f* is a translation inhibitor dose-response curve. Examples are shown in Fig. 11b.

Model variation discussed above can be expanded in a model in which both *r*_min_ and *r*_max_ are varied simultaneously (Fig. 11a). Here, we assume that *r*_min_ increases and *r*_max_ decreases in concert such that they have the same value at *g*_Δr_ = 0 (Fig. 11a). It follows

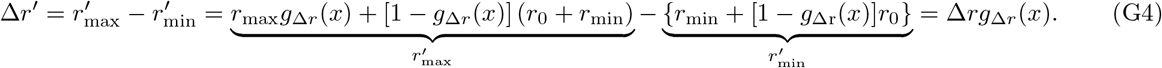

The response parameter *α* is rescaled as above; the ratio *ψ* = IC_50_/IC_50,*F*_ takes a slightly different form:

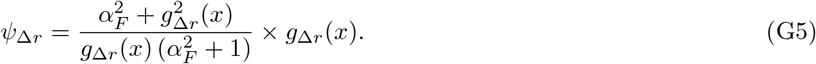

With this at hand, dose-response surfaces can be evaluated (Fig. 11b). Here we note that the both models give the same result if *r*_0_ = Δ*r*, which requires λ_0_ = Δ*rκ_t_*. This is intuitive since the ribosome concentration is already at its maximum and only *r*_min_ can be varied.

## References

[1] C. Walsh, Antibiotics: actions, origins, resistance (ASM Press, Washington DC, 2003).

[2] P. Yeh, M. Hegreness, A. P. Aiden, and R. Kishony, Nat. Rev. Microbiol. 7 (2009).

[3] T. Bollenbach, Curr. Opin. Microbiol. 27, 1 (2015).

[4] S. Loewe, Ergeb. Physiol. 27, 47 (1928).

[5] C. Bliss, Ann. Appl. Biol. 26, 585 (1939).

[6] P. Yeh, A. Tschumi, and R. Kishony, Nat. Genet. 38, 489 (2006).

[7] A. Zimmer, I. Katzir, E. Dekel, A. Mayo, and U. Alon, Proc. Nat. Acad. Sci. USA 113, 10442 (2016).

[8] K. Wood, S. Nishida, E. Sontag, and P. Cluzel, Proc. Nat. Acad. Sci. USA 109, 12254 (2012).

[9] M. Scott, C. Gunderson, E. Mateescu, Z. Zhang, and T. Hwa, Science 330, 1099 (2010).

[10] P. Greulich, M. Scott, M. Evans, and R. Allen, Mol. Syst. Biol. 11, 796 (2015).

[11] M. Scott and T. Hwa, Curr Opin Biotechnol 22, 559 (2011).

[12] B. Kavčič, G. Tkačik, and T. Bollenbach, bioRxiv 10.1101/843920 (2019).

[13] H. Bremer and P. Dennis, in Escherichia coli and Salmonella, edited by F. Neidhardt (ASM Press, Washington DC, 1996).

[14] P. Greulich, J. Doležal, M. Scott, M. Evans, and R. Allen, Phys. Biol. 14 (2017).

[15] B. Davis, Microbiol. Rev. 51, 341 (1987).

[16] J. Elf, K. Nilsson, T. Tenson, and M. Ehrenberg, Phys. Rev. Lett. 97, 258104 (2006).

[17] J. Deris, M. Kim, Z. Zhang, H. Okano, R. Hermsen, A. Groisman, and T. Hwa, Science 342, 1237435 (2013).

[18] B. Kramer and M. Fussenegger, Proc. Nat. Acad. Sci. USA 102, 9517 (2005).

[19] T. Auerbach et al., Proc. Nat. Acad. Sci. USA 107, 1983 (2010).

[20] D. Wilson, Nature Rev. Microbiol. 12, 35 (2014).

[21] M. Belousoff et al., Proc. Nat. Acad. Sci. USA 107, 2717 (2011).

[22] M. Scott, S. Klumpp, E. Mateescu, and T. Hwa, Mol. Syst. Biol. 10, 747 (2014).

[23] M. Rodnina, Cold Spring Harb Perspect Biol 10, a032664 (2018).

[24] J. Elf and M. Ehrenberg, Biophys. J. 88, 132 (2005).

[25] O. Maaløe, Biological Regulation and Development (Plenum, 1979) Chap. Regulation of the Protein-Synthesizing Machinery - Ribosomes, tRNA, Factors, and So On, p. 487x.

[26] T. Durfee et al., J. Bacteriol. 190, 1084 (2008).

[27] R. Cassels, B. Oliva, and D. Knowles, J. Bacteriol. 177, 5161 (1995).

[28] J. Huges and G. Mellows, Biochem. J. 176, 305 (1978).

[29] H. Rode, D. Hanslo, P. Dewet, A. Millar, and S. Cywes, Antimicrob. Agents Chemother. 33, 1358 (1989).

[30] G. Chevereau et al., PLoS Biology 13, e1002299 (2015).

[31] K. Wood, K. Wood, S. Nishida, and P. Cluzel, Cell Rep. 6, 1073 (2014x).

[32] G. Chevereau and T. Bollenbach, Mol. Syst. Biol. 11, 807 (2015).

[33] R. Chait, A. Craney, and R. Kishony, Nature 446, 668 (2007).

[34] R. Lutz and H. Bujard, Nucleic Acids Res. 25, 1203 (1997).

[35] H. Garcia, H. Lee, J. Boedicker, and R. Phillips, Biophys. J. 101, 535 (2011).

[36] M. Steinrück and C. Guet, eLife 6:e25100 (2017).

[37] S. Datta, N. Constantino, and D. Court, Gene 379, 109 (2006).

[38] K. Datsenko and B. Wanner, Proc. Nat. Acad. Sci. USA 96, 6640 (2000).

[39] P. Cherepanov and W. Wackernagel, Gene 158, 9 (1995).

